# Bacterial factors drive the differential targeting of Guanylate Binding Proteins to *Francisella* and *Shigella*

**DOI:** 10.1101/2021.06.16.448779

**Authors:** Stanimira V. Valeva, Fanny Michal, Manon Degabriel, John R. Rohde, Felix Randow, Robert K. Ernst, Brice Lagrange, Thomas Henry

**Author notes:** These authors contributed equally to this work (Fanny Michal, Manon Degabriel).

## Abstract

Guanylate-Binding Proteins (GBPs) are interferon-inducible GTPases that play a key role in cell autonomous responses against intracellular pathogens. Seven GBPs are present in humans. Despite sharing high sequence similarity, subtle differences among GBPs translate into functional divergences that are still largely not understood. A key step for the antimicrobial activity of GBPs towards cytosolic bacteria is the formation of supramolecular GBP complexes on the bacterial surface. Such complexes are formed when GBP1 binds lipopolysaccharide (LPS) from *Shigella* and *Salmonella* and further recruits GBP2, 3, and 4.

Here, we investigated GBPs recruitment on *Francisella novicida*, a professional cytosol-dwelling pathogen with an atypical tetra-acylated LPS. Co-infection experiments demonstrated that GBPs target preferentially *S. flexneri* compared to *F. novicida*. *F. novicida* was coated by GBP1 and GBP2 in human macrophages but escaped targeting by GBP3 and GBP4. GBP1 and GBP2 features that drive recruitment to *F. novicida* were investigated revealing that GBP1 GDPase activity is required to initiate GBP recruitment to *F. novicida* but facultative to target *S. flexneri*. Furthermore, analysis of chimeric GBP2/5 proteins identified a central domain in GBP2 necessary and sufficient to target *F. novicida.* Finally, a *F. novicida* Δ*lpxF* mutant with a penta-acylated lipid A was targeted by GBP3 suggesting that lipid A tetra-acylation contributes to escape from GBP3. Altogether our results indicate that GBPs have different affinity for different bacteria and that the repertoire of GBPs recruited onto cytosolic bacteria is dictated by GBP-intrinsic features and specific bacterial factors, including the structure of the lipid A.

**IMPORTANCE:** Few bacteria have adapted to thrive in the hostile environment of the cell cytosol. As a professional cytosol-dwelling pathogen, *S. flexneri* secretes several effectors to block cytosolic immune effectors, including GBPs. This study illustrates a different approach of adapting to the host cytosol: the stealth strategy developed by *F. novicida*. *F. novicida* bears an atypical hypoacylated LPS, which does not elicit neither TLR4 nor caspase-11 activation. Here, this atypical LPS was shown to promote escape from GBP3 targeting. Furthermore, the lower affinity of GBPs for *F. novicida* allowed to decipher the different domains that govern GBP recruitment to the bacterial surface. This study illustrates the importance of investigating different bacterial models to broaden our understanding of the intricacies of host-pathogen interactions.

## INTRODUCTION

Guanylate-Binding Proteins (GBPs) are interferon-inducible dynamin-like GTPases that play an essential role in host defenses against a large variety of cytosolic pathogens including viruses, intracellular protozoa and bacteria. *GBPs* are present as a multigene family in vertebrates. The *GBP* family exhibits signs of strong evolutionary pressure with gene loss, gene duplication and neofunctionalisation (1–3) indicative of a selective adaptation to pathogens. Eleven GBPs are encoded in mice while seven GBPs are present in humans (4). Human GBPs share a high degree of sequence homology and carry a conserved N-terminal globular GTPase domain followed by a C-terminal helical domain. GTPase activity is required for the antimicrobial activity of GBPs (5–7) and allows GBP dimerization and polymerization (8–10). GTP hydrolysis is well conserved between GBPs. GBP1 is further able to hydrolyze GDP to GMP (11) (Table 1). Additionally, three of the seven GBPs (GBP1, 2 and 5) present a C-terminal CAAX motif and undergo prenylation – i.e. a post-translational addition of a farnesyl or geranylgeranyl lipid group (Table 1). The prenylation allows GBP1, 2, and 5 to be targeted to membranes where they can recruit non-prenylated GBP3/4 (8). Pioneer GBPs recruit downstream GBPs through heterotypic interactions resulting in the formation of supramolecular complexes containing several distinct GBPs (12). GBP5 is particularly targeted to the Golgi apparatus, where it displays anti-viral activity by inhibiting viral glycoprotein maturation (13).

**Table 1.** Properties of macrophage-expressed GBPs

|  | CAAX box | Prenylation | GDP hydrolysis | Closest homologue (s) |
| --- | --- | --- | --- | --- |
| **GBP1** | CTIS | F (15C) | 85% (11) | GBP3 |
| **GBP2** | CNIL | GG (20C) | 5.6% (11) | GBP1 |
| **GBP3** | none | none | N/A | GBP1 |
| **GBP4** | none | none | N/A | GBP7 |
| **GBP5** | CVLL | GG (20C) | 0% (38) | GBP1, 3 |

\*GMP formation calculated from Rajan *et al.* (11) at GMP:GDP ratio for GBP1 = 5.7 and GBP2 = 0.06 F = farnesyl; GG = geranylgeranyl

The antibacterial and anti-parasitic actions of GBPs are well established and most of them are associated with a striking recruitment of GBPs at the pathogen-containing vacuoles (PCV) or directly onto the pathogen if present or released in the cytosol. IFN-γ treatment of *Toxoplasma gondii*-infected cells leads to the recruitment of several thousand of mGBP1/2/3/6 proteins and the formation of a densely packed mGBP coat onto the PCV (12). This recruitment is followed by disruption of the PCV and the ensuing GBP-targeting and lysis of the parasite in the host cytosol. Similarly, GBPs can display antibacterial responses through the targeting of cytosolic bacteria (14–18). These GBP-mediated responses have been particularly well studied with two enterobacteria, *Shigella flexneri* and *Salmonella enterica* serovar *Typhimurium (S. typhimurium)* and have demonstrated a hierarchy in GBP recruitment. Indeed, GBP1 directly binds lipopolysaccharide (LPS) and recruits GBP2, GBP3 and GBP4 to the bacterial surface (15, 19, 20). GBP1 is thus considered as the master GBP orchestrating downstream GBP recruitment.

Assembly of the GBP multimer onto cytosolic bacteria has several consequences. First, as shown for *S. flexneri* and *B. thailandensis*, it can inhibit the formation of actin tails, bacterial motility and cell to cell spread (15, 21, 22). Second, GBPs at the bacterial surface act as a signaling platform to recruit and activate caspase-4, leading to the activation of the non-canonical inflammasome, thus triggering pyroptosis and releasing the pro-inflammatory cytokine interleukin 18 (19, 23). Interestingly, the monitoring of caspase-4 recruitment and activation allowed to ascribe specific functions to the different GBPs. GBP1 acts as the initiator GBP: it is recruited first, exposes bacterial lipid A (the LPS moiety recognized by caspase-4) and elicits both caspase-4 (20) and GBP2/3/4 recruitment. GBP2, 3 and 4 also display redundant or specific roles in caspase-4 recruitment and activation, that are still not fully understood (19, 23). Altogether, these studies suggest that the recruitment of multiple GBPs at the bacterial surface can promote different antibacterial functions. Although highly homologous, growing evidence indicates that subtle differences in GBP sequence and structure may account for functionally relevant divergence. GBP1 was the first GBP to be crystalized and has been extensively characterized (24–28). In contrast, the specific structure-function relationship for other GBPs remains largely unknown.

Although IFN and IFN-inducible proteins, including GBPs, are potent antimicrobial agents against intracellular bacteria, professional cytosol-dwelling bacteria can replicate to very high number in the host cytosol suggesting that they have developed strategies to hide from or actively inhibit GBPs action. Accordingly, *S. flexneri* expresses a Type III secreted effector, IpaH9.8, displaying E3 ubiquitin ligase functions. IpaH9.8-mediated GBP ubiquitination addresses GBPs to the proteasome for proteolytic degradation. Consequently, *S. flexneri* escapes from GBPs-mediated growth restriction (14, 15, 21). *Francisella tularensis,* the agent of tularemia, is another professional cytosolic Gram-negative pathogen that escapes GBP-mediated growth restriction although to different extents depending on the subspecies considered (16, 29). *F. tularensis* can infect a large number of host cells but has a particular tropism for phagocytic cells, including macrophages (30). Following phagocytosis, *F. tularensis* rapidly escapes into the host cytosol using an atypical type IV secretion system (encoded in the Francisella Pathogenicity Island-FPI) (31). *F. tularensis* subsp. *novicida* (hereafter referred to as *F. novicida*) is avirulent in immuno-competent individuals but can infect human cells with a similar life cycle as the highly virulent *F. tularensis* subspecies *tularensis* strains and cause a tularemia-like disease in mice. *F. novicida* has emerged as a model pathogen to study cytosolic immune responses (17, 32–34). *F. novicida* carries an atypical LPS with tetra-acylated lipid A, which enables escape from the host LPS receptors TLR-4 and the murine caspase-11 (33, 35). *F. novicida* LPS can be recognized by caspase-4, in human cells, although it requires one order of magnitude higher concentration than enterobacterial LPS to elicit similar responses (33). In the host cytosol, *F. novicida* is recognized by the AIM2 inflammasome in mice, or the caspase-4 in primary human macrophages (33). Inflammasome activation in mice and in human macrophages (hMDMs) is mediated by GBPs (17, 36). Particularly in hMDMs, GBP2 is recruited to *F. novicida* (33). Still, a comprehensive view of the specific recruitment of GBPs on this cytosolic stealth pathogen is lacking.

In this study, we demonstrate that, in contrast to cytosolic enterobacteria, *F. novicida* escapes GBP3 and GBP4 targeting. Targeting of GBP3 was partially restored in a Δ*lpxF* mutant, presenting a penta-acylated lipid A. Furthermore, through co-infection experiments we revealed that GBP1 targets preferentially *S. flexneri* compared to *F. novicida*. Finally, GBP chimeras-based structure-function analyses identified the specific domains in GBP1 and GBP2 driving recruitment to *F. novicida*. These analyses revealed distinct GBP features required to target to *F. novicida* but facultative for recruitment to *S. flexneri.* Altogether, our results suggest that, in contrast to the prevailing model, GBP recruitment downstream of GBP1 is not only driven by heterotypic GBP interactions but results from multiple GBP-intrinsic features. Further, GBP targeting is controlled by bacterial factors including but not restricted to, lipid A acylation levels.

## RESULTS

### *F. novicida* specifically escapes GBP3-4 targeting

To study the specific recruitment of individual human GBP to *F. novicida*, we generated stable human monocyte/macrophage U937 cell lines constitutively expressing HA-tagged GBP 1-5 (Fig. S1A). GBP6 and 7 were not studied since they are not expressed at substantial level in monocyte-derived macrophages (33). As observed in primary human macrophages, endogenous GBP1-5 were highly induced in U937 macrophages upon IFNγ treatment (Fig. S1B). IFNγ-primed, PMA-differentiated U937 macrophages were infected with *F. novicida* and imaged 7 h post-infection to assess specific GBP recruitment. GBP1 and GBP2 were targeted to a subset of bacteria (Fig. 1A, C) and intimately colocalized with *F. novicida* in ≈25% of infected cells. Surprisingly, targeting of GBP3 or GBP4 to *F. novicida* could not be observed. The absence of GBP3/4 recruitment onto *F. novicida* contrasted with previous studies using other Gram-negative pathogens. Indeed, GBP1, 2, 3 and 4 are recruited to *S. flexneri* and *S. enterica* serovar Typhimurium *(S. typhimurium)* in HeLa cells (15, 19, 23). Importantly, GBP3 and GBP4 (and GBP1/2) were recruited to *S. flexneri* ΔipaH9.8 (hereafter referred to as *S. flexneri*) as early as 3 h p.i. in infected U937 macrophages (Fig. 1B, D), demonstrating the functionality of these GBPs in our experimental system. As expected, GBP5, which is recruited to neither *S. typhimurium* nor *S. flexneri,* was not recruited to *F. novicida* either.

**FIG 1.**
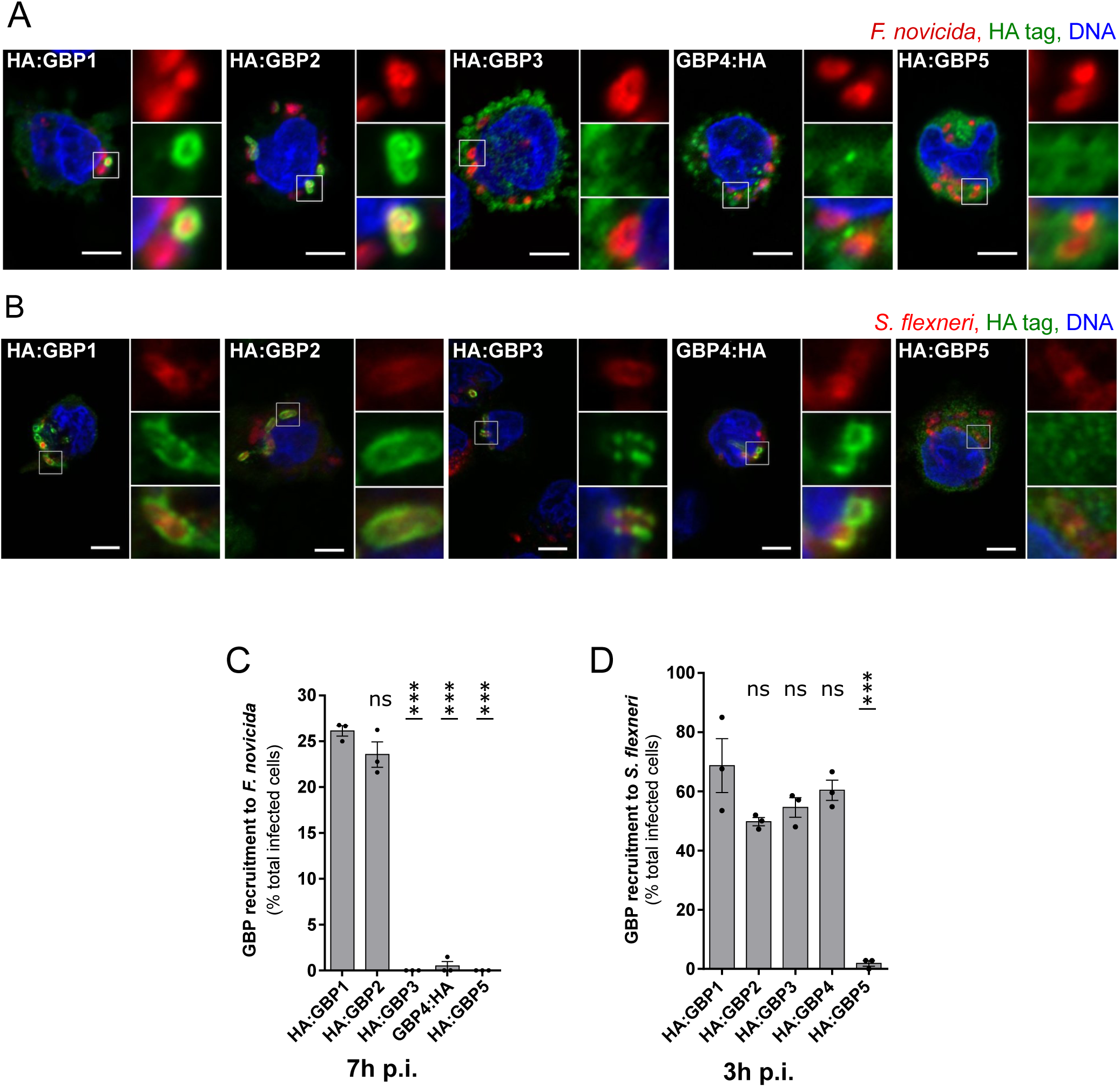
*F. novicida* and *S. flexneri* are targeted by a different repertoire of GBPs. IFNγ-treated, HA:GBP-expressing, U937 macrophages were infected with *F. novicida* (A, C) or *S. flexneri* ΔipaH9.8 (B, D) at the indicated time post-infection (p.i.). (A) and (B) show representative images, scale bar 5 µm and 3X zoom on the right panels. (C) and (D) GBP recruitment was quantified as the percent of infected cells in which GBPs are targeted to bacteria. Each point represents the value from one experiment with 50-100 infected cells counted per experiment. The bar represents the mean +/- SEM of three independent experiments. ANOVA with Dunnett’s analysis was performed in comparison to GBP1 recruitment frequency: ***, p < 0.001; ns, not significant.

These findings show that the repertoire of GBPs recruited to *F. novicida* and to *S. flexneri* differs, suggesting that *F. novicida* escapes targeting by GBP3 and 4.

### Recruitment of GBP2 to *F. novicida* depends on GBP1

Studies with *S. typhimurium* and *S. flexneri* have established that GBP1 initiates recruitment of GBP2, 3 and 4 to the bacterial surface (15, 19, 23). As the above results demonstrated differences in GBP targeting between *F. novicida* and the previously studied enterobacteria, we examined the hierarchy of GBP1 and GBP2 targeting to *F. novicida*.

HA:GBP1 and HA:GBP2 were expressed in *GBP*^KO^ U937 cells (Fig. S2A, B) and their recruitment to *F. novicida* was scored at 7 h post-infection. Individual *GBP2-5* knock-out did not affect the frequency of HA:GBP1 recruitment to *F. novicida* suggesting that GBP1 is recruited independently of other GBPs (Fig. 2A, B). Accordingly, in the absence of IFNγ (and hence of endogenous GBPs), ectopically expressed HA:GBP1 was recruited to *F. novicida* (Fig. S2C, D). In contrast, HA:GBP2 was not recruited to *F. novicida* in *GBP1*^KO^ cells (Fig. 2A, C), nor in the absence of IFNγ (Fig. S2C, D). Recruitment of HA:GBP2 was not affected by invalidation of *GBP3, 4* or *5*. Thus, as previously reported for *S. flexneri* (15, 19, 23) and *S. typhimurium* (19), GBP1 is recruited to *F. novicida* independently of other GBPs and of other IFNγ-induced factors whereas GBP2 recruitment requires expression of GBP1.

**FIG 2.**
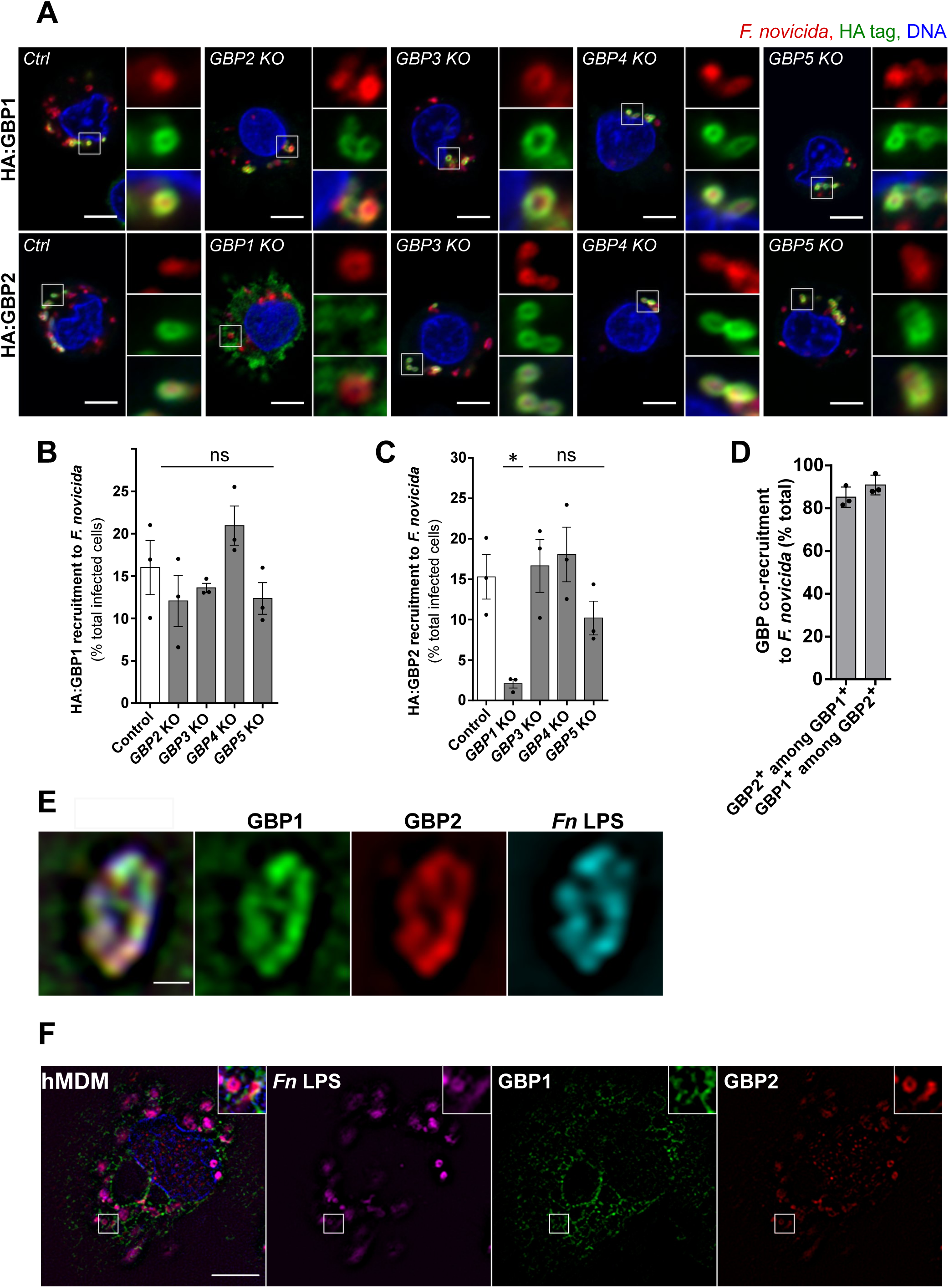
GBP2 is recruited in a GBP1-dependent manner to *F. novicida*. IFNγ-treated U937 (A-E) or monocyte-derived macrophages (F) were infected with *F. novicida* for 7 h (A-E) or 10 h (F). (A) Representative images of *GBP^KO^* or control U937 cells stably expressing HA:GBP1 (top panels) or HA-GBP2 (lower panels) are shown with scale bar 5µm and 3X zoom. (B, C) HA:GBP recruitment was expressed as the percentage of infected cells presenting GBP-bacteria colocalization. (D) Endogenous GBP co-recruitment to *F. novicida* measured as the percent of GBP2-positive bacteria among the GBP1-positive and vice-versa. (E, F) Structured illumination microscopy of endogenous GBP localization to *F. novicida* in U937 cells (E) or human monocyte-derived macrophages (hMDMs) (F), scale bar 0.5µm in (E) and 5 µm in (F). Data information (B-D): Each point indicates the value of one experiment with 50-100 infected cells analyzed. The bar represents the mean +/-SEM of 3 independent experiments. ANOVA with Dunnett’s analysis was performed in comparison to recruitment frequency in control cells: *, p < 0.05; ns, not significant.

Because of the high homology between the GBPs (Table 1), few tools exist for studying the specific recruitment of endogenous GBPs. We validated antibodies to specifically immunolabel GBP1 and GBP2 (Fig. S2E). Using these antibodies, we investigated the co-recruitment of GBP1 and GBP2 to *F. novicida* in wild-type cells. Whenever one GBP (GBP1 or GBP2) was targeted to a bacterium, the other GBP was present on the same bacterium in more than 85% of the cases (Fig. 2C). High-resolution images of structured illumination microscopy (SIM) demonstrated intimate colocalization between endogenous GBP1, GBP2, and the bacterial LPS in U937 macrophages, and in primary human macrophages (Fig. 2 E, F). Therefore, once recruited, GBP1 consistently recruits GBP2 onto *F. novicida* where both proteins tightly colocalize with LPS on the bacterial surface.

### GBP2 CAAX box increases targeting to *F. novicida* compared to GBP1 CAAX box

GBP1 and GBP2 undergo a post-translational addition of a lipid prenyl group (8). This prenylation is guided by a C terminal CAAX motif, which facilitates attachment of proteins to cell membranes (Table 1). Prenylation is required for GBP1 to target *S. flexneri* (19–21) and *S. typhimurium*, (7, 19) and for GBP-dependent inflammasome activity. (7) Deletion of the GBP1 CAAX box (Fig S3. A) also abolished GBP1 recruitment to *F. novicida* (Fig 3.A) Similarly, deletion of GBP2 CAAX box abrogated recruitment to *F. novicida* (Fig 3.B, D), and to *S. flexneri* (Fig 3.C, D). These data reveal that prenylation is necessary for GBP1 and GBP2 targeting to *F. novicida*.

**FIG 3.**
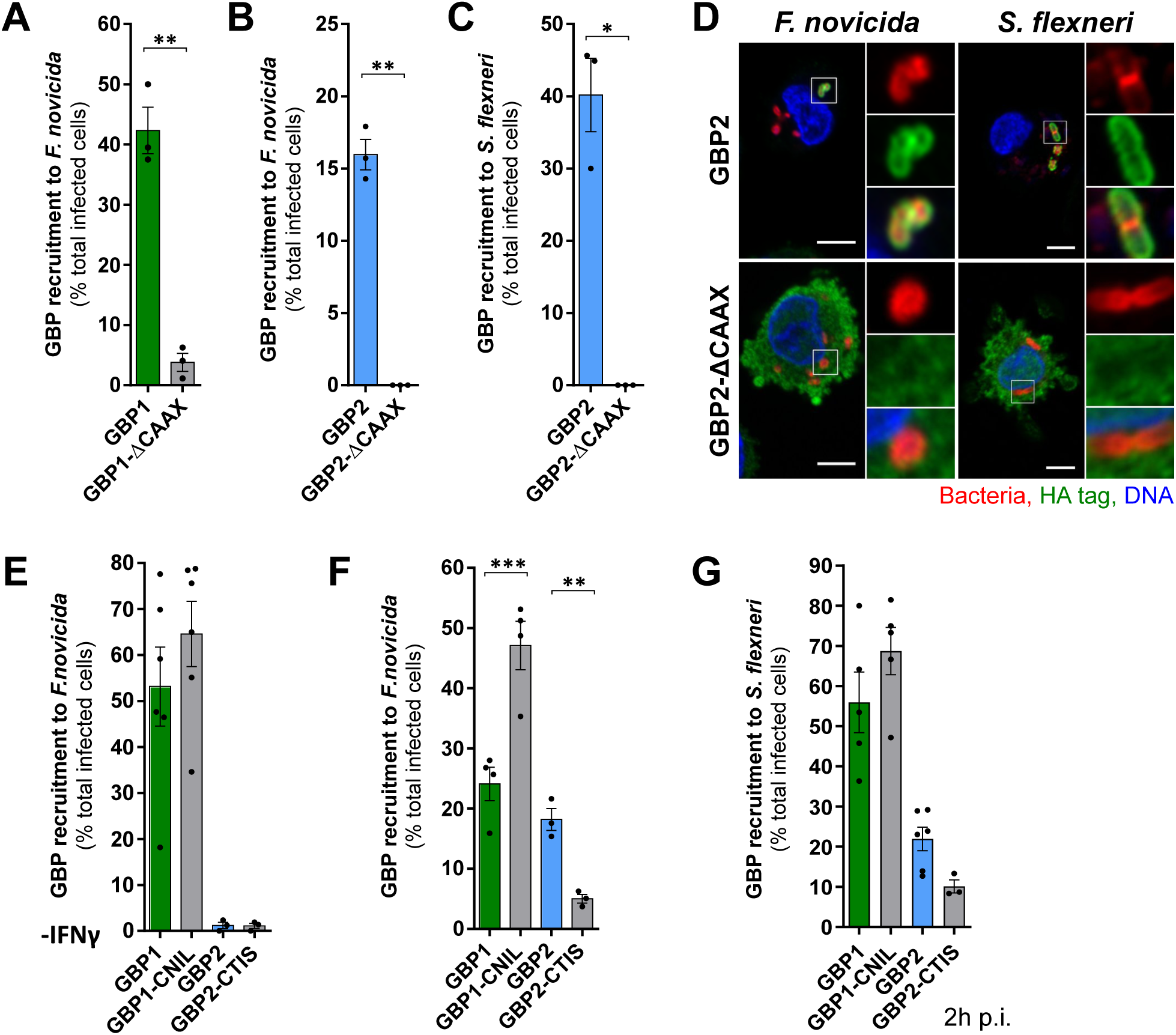
GBP2 CAAX box increases GBP recruitment to *F. novicida.* U937 macrophages treated (A-D, F, G) or not (E) with IFNγ, were infected with *F. novicida* for 7 h (A, B, D-F) or *S. flexneri* ΔipaH9.8 for 2 (G) to 3 h (C). (D) Representative images with scale bar 5µm and 3X (*F. novicida*) or 2X (*S. flexneri*) zoom on the right panels are shown. (A-C, E-G) GBP recruitment was scored as the percentage of infected cells with GBP-bacteria colocalization. Each point corresponds to the value from one experiment with 50-100 infected cells analyzed. The bar represents the mean +/- SEM of three independent experiments. Two-tailed t test with Welch’s correction (A-C) or ANOVA with Sidak’s (E-G) analysis was performed: *, p <0.05; **, p < 0.01; ***, p < 0.001.

Interestingly, the lipid moiety of GBP2 is a geranylgeranyl lipid that consists of a 20 carbon chain whereas GBP1 lipid moiety is a farnesyl, which is a shorter 15 carbon chain (Table 1). The type of prenylation is dictated by the sequence of the CAAX box (8, 37). We thus wondered whether the difference in prenylation between GBP1 and 2 might play a role in bacteria targeting. GBP1 and GBP2 CAAX motives were swapped to generate cell lines expressing HA:GBP1-CNIL or GBP2-CTIS (Fig. S3A). In the absence of IFNγ, both GBP1 and GBP1-CNIL were recruited to *F. novicida* while GBP2 and GBP2-CTIS were not (Fig. 3E). Therefore, GBP farnesylation is not sufficient to initiate GBP recruitment. Remarkably, HA:GBP1-CNIL was consistently recruited at higher rates than HA:GBP1 to *F. novicida*. The increased recruitment of GBP1-CNIL was even more striking upon IFNγ treatment. Conversely, GBP2-CTIS recruitment was significantly lower than that of HA:GBP2 (Fig. 3F, S3C) to *F. novicida*. A similar trend was observed with GBP targeting to *S. flexneri* (Fig. 3G), albeit not statistically significant. Additionally, GBP1-CNIL-expressing cells responded to *F. novicida* infection with faster and higher cell death rates than the other cell lines (Fig. S3D), suggesting that the increased recruitment has functional consequences on inflammasome-mediated cell death.

Altogether, these results reveal that GBP prenylation is required to target *F. novicida.* While the specificity of GBP prenyl chain does not drive GBP recruitment hierarchy, GBP prenylation type may control GBP recruitment efficiency. Indeed, the CNIL CAAX box associated with geranylgeranylation boosts GBP targeting to *F. novicida* compared to the CTIS CAAX box associated with farnesylation.

### The central region of GBP2 controls its recruitment to *F. novicida*

Besides GBP1 and GBP2, GBP5 is the only other GBP presenting a CAAX box. Yet, GBP5 is not recruited to bacteria (Fig. 1) (15, 19, 23). GBP5 is modified by geranylgeranylation similarly to GBP2 (8). As expected, replacing the CAAX box of GBP2 with that of GBP5 (GBP2-CVLL, Fig. S4A) did not significantly alter recruitment. Likewise, GBP5 carrying the CAAX box of GBP2 (GBP5-CNIL) did not colocalize with *F. novicida* (Fig. 4A). Thus, although necessary for GBP1/2 targeting to *F. novicida*, GBP prenylation by itself is not sufficient for a GBP to be targeted to bacteria indicating that additional domains govern the selective recruitment of GBP2 to *F. novicida*.

**FIG 4.**
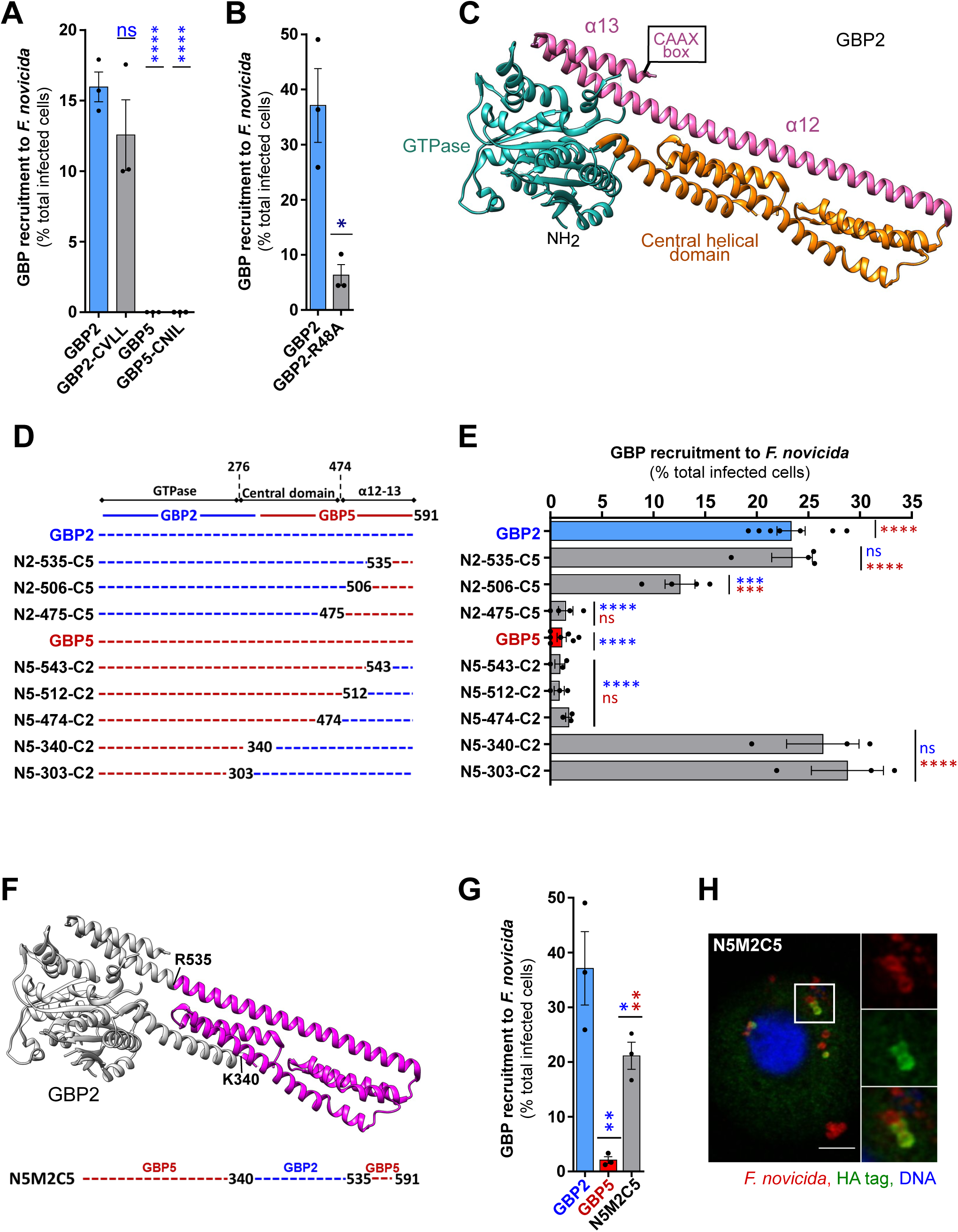
The central domain of GBP2 controls GBP2 recruitment to *F. novicida*. (A, B, E, G, H) IFNγ-treated, U937 macrophages were infected with *F. novicida*. GBP recruitment was quantified as the percentage of infected cells with *F. novicida*-HA-GBP colocalization. (C) GBP2 structure is presented with coloring corresponding to the studied chimera. (D) GBP2-5 or GBP5-2 chimera stably expressed in U937 cells are schematically shown. (F) The GBP2 region identified as necessary and sufficient to promote GBP2 recruitment is presented with the corresponding schematic GBP5-GBP2-GBP5 chimera, underneath. (H) Representative image is shown with scale bar 5 µm and 2X zoom on the right panels. Two-tailed t test with Welch’s correction (B) or ANOVA with Dunnett’s (A, E, G) analyses were performed: *, p <0.05; **, p< 0.01; ***, p <0.001; ****, p <0.0001; ns, not significant. Blue and red star/writing indicates the result of the statistical comparison with GBP2 or GBP5, respectively.

The GTPase activity of GBP2 was also required to target *F. novicida*. Indeed, a GTPase null mutant GBP2-R48A failed to localize to the bacteria (Fig. 4B, S4B-C). However, GBP5 is also capable of GTP hydrolysis (38) and the catalytic residues, highly conserved in the dynamin superfamily (39), are identical in GBP2 and GBP5 (Fig. S4K).

The crystal structures of GBP2 and GBP5 were solved recently (40). GBP2 and GBP5, similarly to GBP1, contain a globular GTPase domain at the N-terminus, followed by an elongated helical region, which ends on a hairpin-like C-terminus with the CAAX motif at the very end (Fig. 4C). GBP2 and GBP5 also share a high similarity, with the most divergence localized in the C-terminal α12 and α13 helices (Fig. S4K). Owing to the similarity between both proteins, multiple GBP2-GBP5 and GBP5-GBP2 chimeras were generated and stably expressed in U937 cells (Fig. 4D, Fig. S4D). The chimeras were evaluated for gain or loss of targeting to *F. novicida* to pinpoint the specific GBP2 domain driving recruitment. Chimera N2-535-C5, consisting of GBP2 up to residue R535, followed by the GBP5 C-terminus, was recruited to *F. novicida* similarly to GBP2 (Fig. 4E). Thus, contrary to our expectations, the targeting specificity of GBP2 is not driven by the most C-terminal part of GBP2 (535-end containing α13 and 1/3 of α12). Chimera N2-506-C5 presented a 2-fold decrease in recruitment to *F. novicida* compared to GBP2 while chimera N2-475-C5 was not recruited at all. These observations indicate that recruitment to *F. novicida* gradually decreases as GBP2 C-terminus is replaced by GBP5 between residues Q475 and R535, establishing that this region is required for GBP2 recruitment. However, the presence of this GBP2 Q475-R535 was not sufficient to induce the recruitment of the corresponding GBP5-GBP2 chimera (termed N5-474-C2) (Fig. 4F). Further addition of GBP2 residues K340-L474 generated a chimera (termed N5-340-C2) gaining the full ability to be recruited to *F. novicida*.

These experiments revealed two neighboring regions (K340-L474 and Q475-R535) in the central part of GBP2 which are necessary for bacteria targeting in the context of GBP5-GBP2 and GBP2-GBP5 chimeras, respectively. To assess whether the GBP2 K340-R535 region would be sufficient to drive the recruitment of a prenylated GBP to *F. novicida*, we generated cells stably expressing a chimera with the central domain of GBP2 (340-535) in a GBP5 background (chimera N5M2C5, Fig. 4G). This three-part chimera, although only faintly expressed (Fig. S4E), was indeed recruited to *F. novicida* (Fig. 4H, I) thus identifying the central domain of GBP2 (340-535) as necessary and sufficient in the context of GBP5 to drive recruitment to *F. novicida*.

A unique feature of GBP5 is its localization in the Golgi apparatus (41, 42). We thus wondered whether the Golgi apparatus localization of GBP5 could be responsible for the lack of recruitment to cytosolic *F. novicida*. We first calculated Golgi enrichment ratios for all GBP2/5 chimeras (Fig. S4F-I). Increasing the proportion of GBP5 sequence in the C-terminus gradually increased Golgi apparatus localization in GBP2-GBP5 chimeras. Conversely, an increase in the C-terminal GBP2 proportion paralleled a decrease in Golgi apparatus localization. These results indicate that the central helical domain and the α12-α13 region contribute to GBP5 localization at the Golgi apparatus. Yet, in contrast to recruitment to *F. novicida*, we could not delineate a specific Golgi-targeting domain. We thus analyzed all the GBP2/5 chimeras to assess whether the Golgi apparatus localization was inversely correlated with recruitment to *F. novicida*. No correlation could be observed (Fig. S4J) suggesting that Golgi apparatus localization and recruitment to bacteria are independent GBP features.

Altogether, the chimera recruitment assays uncovered an essential role of the central region of GBP2 (340-535) in *F. novicida* targeting. These results also indicate that the corresponding region in GBP5 diverges from GBP2 to a degree that impedes recruitment of GBP5 to *F. novicida,* independently of GBP5 Golgi localization.

### GMP formation by GBP1 is required for recruitment to *F. novicida* but not to *S. flexneri*

As described above (Fig. 2), GBP2 recruitment to *F. novicida* is governed by the initial GBP1 recruitment, while GBP1 is targeted to bacteria independently of IFNγ-induced factors. To identify the GBP1-specific domains driving the initial recruitment to *F. novicida,* GBP1-GBP2 and GBP2-GBP1 chimeric cell lines were generated (Fig. 5A, Fig. S5A).

**FIG 5.**
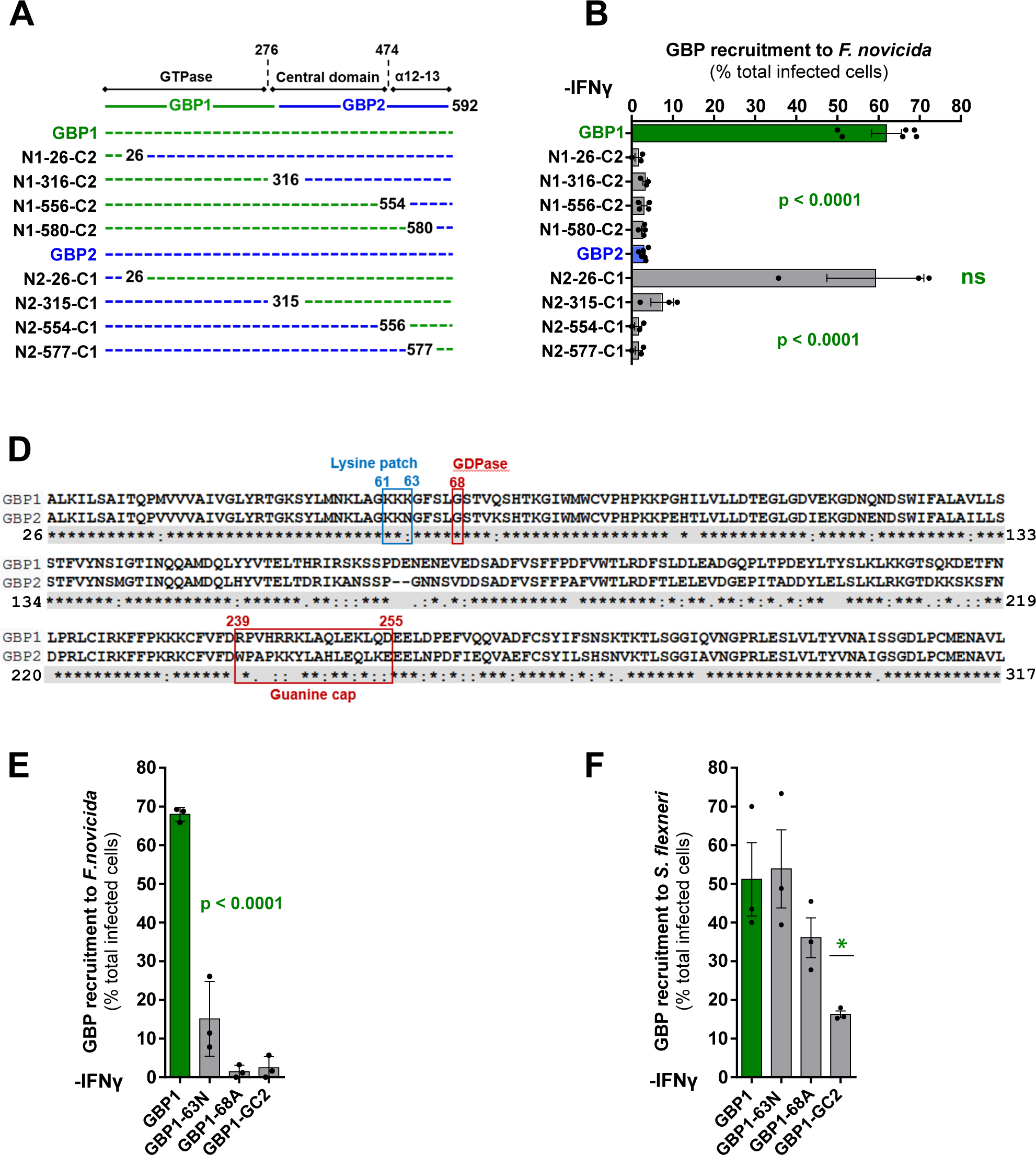
GMP formation by GBP1 is specifically required for IFNγ-independent recruitment to *F. novicida.* (A) GBP1-2 or GBP2-1 chimera are schematically shown. (B, E, F) U937 macrophages were infected with *F. novicida* for 7 h (B, E) or *S. flexneri* ΔipaH9.8 for 2 h (F). Recruitment was calculated as the percentage of infected cells with HA-GBP-bacteria colocalization. (D) Clustal alignment of GBP1 and GBP2 GTPase domains with a highlight on the studied domain/residues. (B, E, F) Each point corresponds to the value from one experiment with 50-100 infected cells analyzed. The bar represents the mean +/- SEM of three to six independent experiments. ANOVA with Sidak’s (B) or Dunnett’s (D, E) analysis was performed in comparison with GBP1 recruitment: *, p <0.05; ns, not significant.

Several studies pinpoint a unique polybasic patch (RRR_584-586_) in the GBP1 C-terminus that controls GBP1 targeting to *S. flexneri* and *S. typhimurium* (1, 21). This CAAX-neighboring sequence is absent in GBP2 and in other human GBPs. The N1-580-C2 chimera (lacking the triple R patch) was not recruited to *S. flexneri* (Fig. S5B) in the absence of IFNγ priming. Conversely, the presence of GBP1 14 last amino-acid residues (including the triple R patch) in the GBP2-1 chimera N2-577-C1 was sufficient to drive its IFNγ-independent recruitment to *S. flexneri*. Similarly, in the absence of IFNγ, none of the GBP2 C-terminal chimeras were recruited to *F. novicida* strongly suggesting that the triple arginine patch also controls *F. novicida* targeting (Fig. 5B, Fig. S5E). However, N2-577-C1 chimera was not recruited to *F. novicida* either. This result indicates that, in contrast to *S. flexneri*, the last 14 amino-acid residues of GBP1 (Q577-end) are not sufficient to promote IFNγ-independent GBP recruitment. Further addition of GBP1 central domain and α12-13 region residues (A315-I576) did not induce IFNγ-independent recruitment of the corresponding GBP2-GBP1 chimeras (N2-551-C1 or N2-315-C1). A GBP1 phenotype was restored in chimera N2-26-C1 which additionally carries the Q26-L316 region of GBP1, corresponding to the globular GTPase domain. Importantly, in the presence of IFNγ, all the above chimeras could be recruited to *F. novicida* thus confirming their functionality in terms of recruitment (Fig. S5C, D).

The GTPase domains of GBP1 and GBP2 present 80% identity and 90% similarity (Fig. 5D). However, the chimera recruitment assays point to a functional difference in the GTPase domains of GBP1 and GBP2 in terms of their ability to specifically initiate recruitment to *F. novicida*. Recently, Santos *et al.* identified several positively charged patches on the surface of GBP1, of which KKK_61-63_, present in the GTPase domain, was required for GBP1 recruitment to *S. typhimurium* (19). GBP2 carries only two lysine residues (KKN_61-63_) at the corresponding location, we thus wondered whether a K63N mutation would influence GBP1 recruitment to *F. novicida*. In addition, GBP1 is the only GBP known to efficiently hydrolyze GDP to GMP (Table 1, (11)). Xavier *et al*. (43) recently identified a GBP1 mutation (G68A) that blocked GDP hydrolysis while leaving GTP hydrolysis intact. The residue in this position is identical in GBP2 (Fig. 5D). However, Rajan *et al*. proposed that GMP formation of GBP1 is due to a “guanine cap” loop, situated between R239 and D255, which stabilizes GDP in the GBP1 catalytic pocket. The guanine cap is also present in GBP2 but, it differs in its tertiary structure and does not promotes GMP formation (11). We thus wanted to determine whether GBP1 GDPase activity may promote its recruitment to *F. novicida*.

To assess the above two hypotheses, cell lines were created to express GBP1 mutated in K63, G68 or a GBP1 variant carrying GBP2 guanine cap (GBP1-GC2) (Fig. S5F). The mutated proteins were functional for recruitment to *F. novicida* upon IFNγ treatment (Fig. S5G). The K63N mutation dramatically reduced GBP1 targeting to *F. novicida* in the absence of IFNγ (Fig. 5E). Thus, the KKK_61-63_ lysine patch contributes to initiating GBP1 recruitment to *F. novicida* and the loss of a single lysine residue is sufficient to strongly limit the ability of the resulting GBP1 mutant to target *F. novicida*. Mutation of the G68 residue in GBP1 completely abolished recruitment to *F. novicida* in the absence of IFNγ. GBP1-GC2 was not recruited to *F. novicida* without IFNγ either. These results strongly suggest that GMP formation by GBP1 is essential for initiating GBP1 recruitment to *F. novicida*. Surprisingly, the K63N mutation did not affect IFNγ-independent GBP1 recruitment to *S. flexneri* (Fig. 5F). The G68A GBP1 mutant was also recruited to *S. flexneri* similarly to GBP1. Replacement of the GBP1 guanine cap with that of GBP2 resulted in a statistically significant decrease in *S. flexneri* targeting. Yet, in 15% of infected cells, GBP1-GC2 chimera was robustly targeted to *S. flexneri* (Fig. S5H) indicating that, in contrast to *F. novicida* targeting, GMP formation by GBP1 is not necessary for recruitment to *S. flexneri*.

Overall, these findings establish a requirement for GMP formation by GBP1 in order to initiate recruitment to *F. novicida*. Furthermore, these results indicate that not only *S. flexneri* and *F. novicida* differ in the repertoire of GBPs targeted to their surface (Fig. 1) but also in the requirement of GBP1 motifs/activity to target the bacterial surfaces.

### Tetra-acylation of *F. novicida* LPS limits GBP recruitment

Suppression of GBP3/4 recruitment might be due to either an active process (e.g. implicating a T6SS-secreted effector) or a lack of recognition (*F. novicida* being a stealth pathogen (44)). The role of the T6SS could not be directly tested since a ΔFPI mutant (lacking the T6SS) does not escape into the cytosol and thus fails to recruit GBPs (33). Treatment of *F. novicida*-infected macrophages with chloramphenicol, an antibiotic blocking protein neosynthesis, did not promote GBP3 recruitment to *F. novicida* (Fig. S6A) suggesting that the absence of GBP3 recruitment onto *F. novicida* is not due to active inhibition.

To further evaluate whether *F. novicida* could actively block GBP3 recruitment via secreted proteins, HA:GBP-expressing macrophages were co-infected with *F. novicida* and *S. flexneri*. The cells were first infected with *F. novicida* and 4 h later with *S. flexneri* until 7 h total to ensure optimal GBP recruitment rates for both species. In co-infected cells, GBPs 1-4 were robustly recruited to *S. flexneri* (Fig. 6A). Similar results were obtained when cells when inoculated with *F. novicida* and *S. flexneri* at the same time (Fig. S6B). Likewise, co-infection of primary human macrophages with *F. novicida* did not suppress the targeting of endogenous GBP2 to *S. flexneri* (Fig 6B). Therefore, the mechanism allowing *F. novicida* to escape GBP3/4 targeting does not act in trans on *S. flexneri* but is restricted to the bacterium.

**FIG 6.**
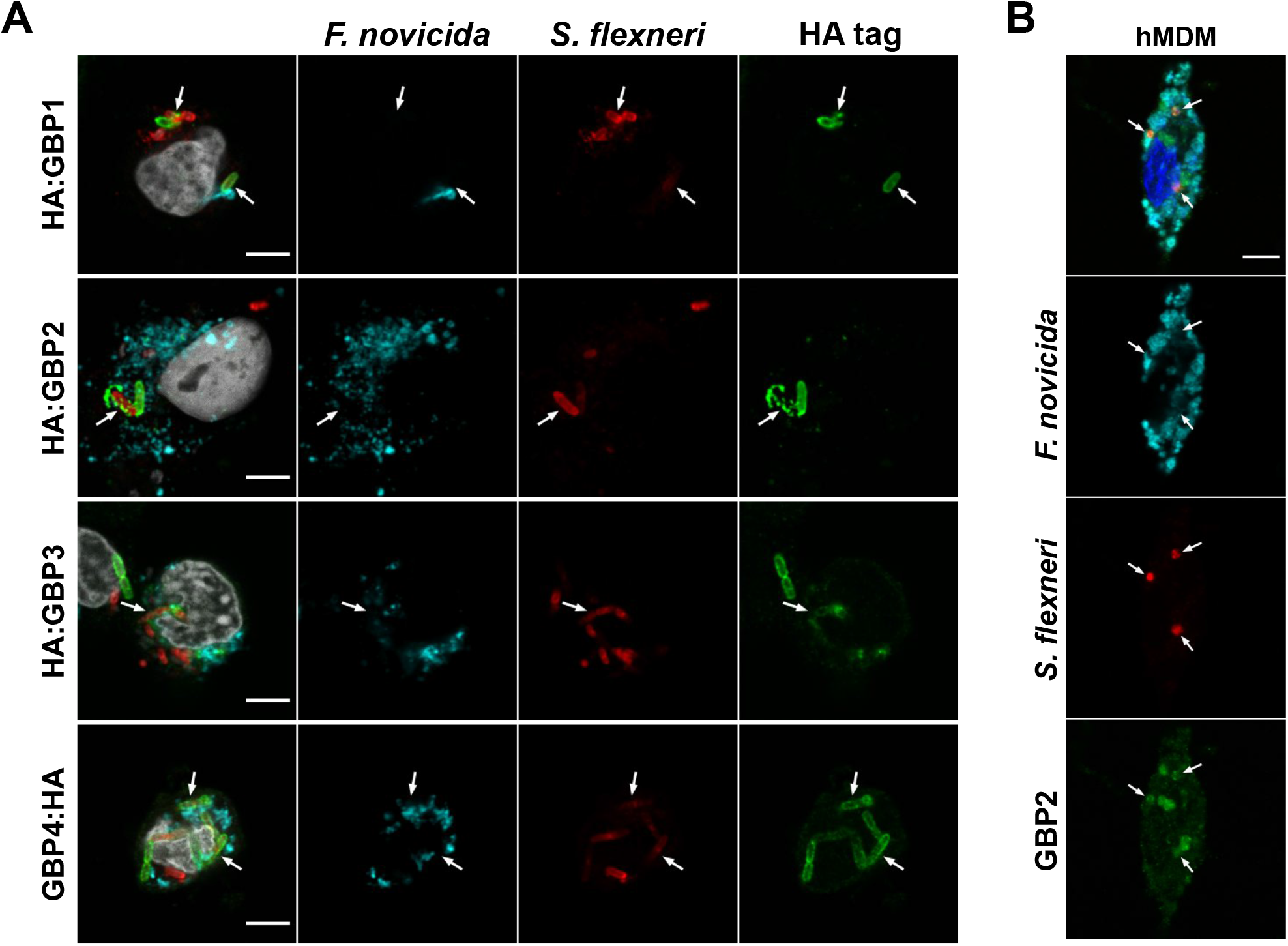
GBPs are selectively recruited to *S. flexneri* in *F. novicida*-co-infected cells. IFNγ-primed U937 macrophages (A) or human monocyte-derived macrophages (hMDMs, B) were infected with *F. novicida* at MOI 50 and then 4 h later with *S. flexneri* ΔipaH9.8 at MOI 50 for 3 h (A) to 4 h (B). Representative images are shown. The arrows indicate GBP recruitment to *S. flexneri*. Scale bar, 5 µm.

Curiously, while GBP1 and GBP2 recruitment could easily be observed in *F. novicida*-infected cells, we could not find any GBP1 or GBP2 recruitment to *F. novicida* in co-infected cells (Fig. 6A). This observation was true for all co-infection experiments, regardless of the time of infection (Fig S6B) and was validated in primary human macrophages (Fig. 6B). The lack of recruitment onto *F. novicida* was restricted to co-infected cells since GBP1/2 recruitment on *F. novicida* was detected in bystander cells infected only by *F. novicida* (Fig. S6C). These results suggest that in co-infected cells, GBP1 and GBP2 target preferentially *S. flexneri* alluding to lower affinity or avidity of GBP1 and 2 for the *F. novicida* envelope than for the one of *S. flexneri*.

The above results rule against an active mechanism used by *F. novicida* to avoid GBP3 and GBP4 targeting. A key feature of *F. novicida* is the atypical LPS containing a tetra-acylated lipid A, which has been associated with immune evasion (33, 45, 46). To explore whether lipid A tetra-acylation plays a role in the escape of *F. novicida* from GBP3/4 recruitment, we infected U937 macrophages with a *F. novicida* Δ*lpxF* mutant. LpxF removes the phosphate in position 4’ of the lipid A. Its absence results in a penta-acylated lipid A (Fig. 7A) due to steric hindrance that blocks the 3’ deacylase, LpxR (35). The Δ*lpxF* mutant had a lower ability than the WT strain to rupture its phagosome (Fig. S7A), which was reflected in the lower rates of GBP1 recruitment (Fig. S7B). Nevertheless, GBP1 recruitment was robust and clearly visible (Fig. S7C). We thus used endogenous GBP1 as a marker of cytosolic bacteria and analyzed HA:GBP3 and HA:GBP4 recruitment. We could not observe a robust HA:GBP3 or HA:GBP4 recruitment at the surface of the GBP1^+^ Δ*lpxF* mutant strain as can be seen in *S. flexneri*-infected cells (Fig. 1B) or for HA-GBP1/2 on the surface of *F. novicida* (i.e. clear accumulation at the bacterial surface associated with a depletion of the cytosolic diffuse staining). Yet, we consistently noticed a discrete recruitment of HA-GBP3 on GBP1^+^ Δ*lpxF* mutant strain (Fig. 7C). These observations were quantified using confocal images by scoring the enrichment of HA:GBP3 at the surface of GBP1^+^ bacteria (Fig. S7D). HA:GBP3 was significantly more enriched on the Δ*lpxF* mutant than on the WT strain (Fig. 7B, n = 26, p < 0.0001). Images corresponding to the highest and to the mean HA-GBP3 enrichment ratio are presented in Fig. 7C for the WT and Δ*lpxF* mutant strain. The images clearly illustrate a specific, although low, recruitment of HA-GBP3 on the Δ*lpxF* mutant strain that is not observed on WT *F. novicida*. No visible (Fig. S7F) nor quantifiable (Fig. S7E) enrichment of GBP4-HA could be detected on Δ*lpxF* mutant strain demonstrating the specificity of GBP3 localization to penta-acylated Δ*lpxF F. novicida*.

**FIG 7.**
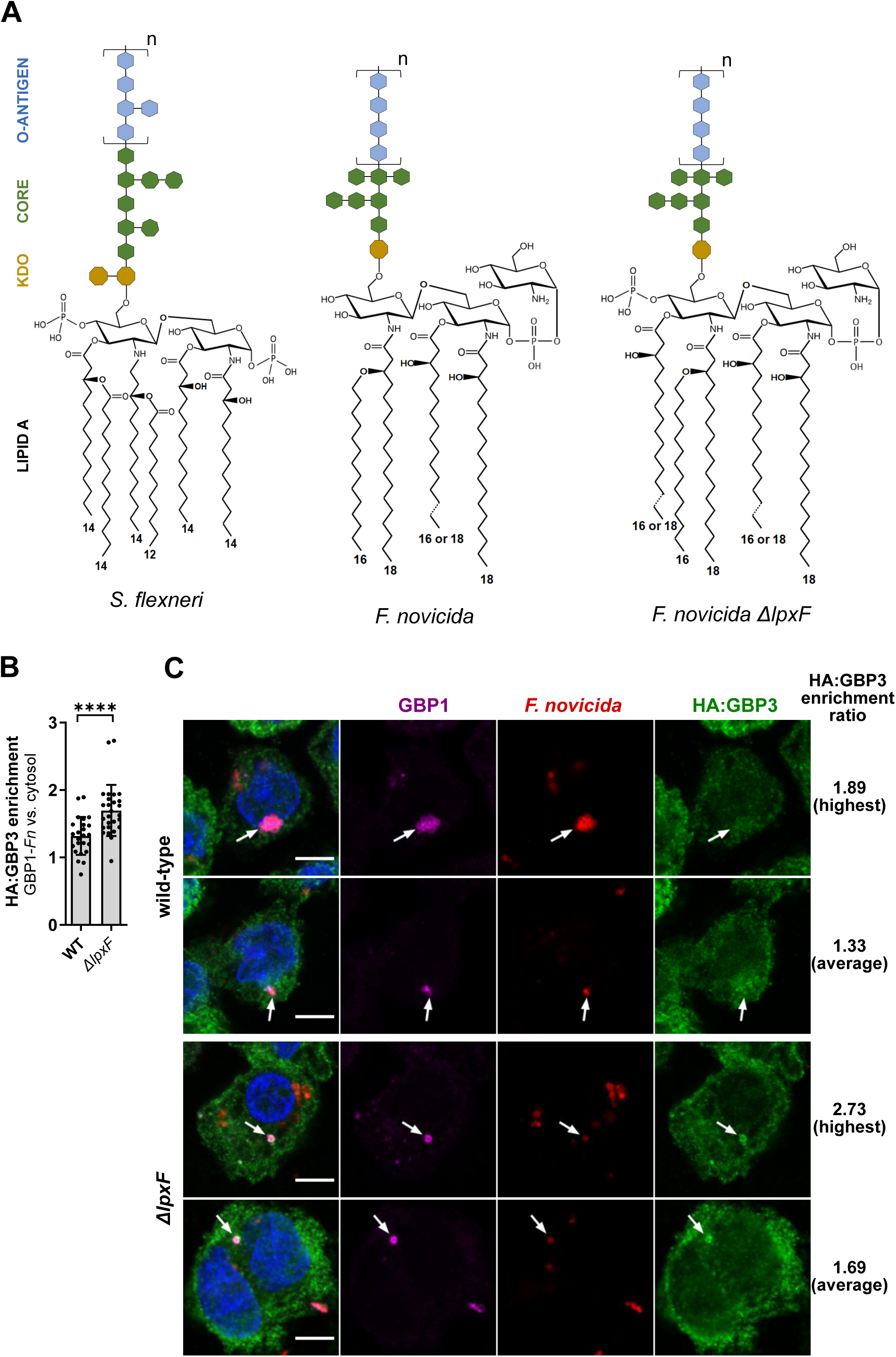
LPS penta-acylation promotes GBP3 targeting to *F. novicida* (A) Structure of hexa-acylated *S. flexneri*, tetra-acylated *F. novicida* and penta-acylated *F. novicida* Δ*lpxF* LPS. (B, C) IFNγ-treated, U937 HA:GBP3-expressing macrophages were infected with *F. novicida* for 7 h. HA:GBP3 enrichment was calculated as explained in Fig. S7B. (B) Each point represents the value of HA:GBP3 enrichment on one individual GBP1-*F. novicida* colocalization area. The bar represents the mean +/- SEM of 26 events originating from 4 independent experiments. Two-tailed Mann Whitney analysis was performed: ****, p<0.0001. (C) Images with the highest or the average GBP3 enrichment value are shown as indicated. Scale bar, 5µm.

Altogether, these results indicate that *F. novicida* escapes GBP3 recognition owing to its atypical tetra-acylated LPS. However, GBP3 targeting to the Δ*lpxF* mutant was not as pronounced as a “typical” GBP recruitment seen on other bacteria or seen with GBP1/2 onto *F. novicida*. Thus, additional prokaryotic factors likely constrain GBP3 (and to an even greater extent, GBP4) targeting to bacteria.

## DISCUSSION

Professional cytosol-dwelling bacteria either hide from or actively inhibit cell autonomous responses to thrive in the host cytosol. Here, we observed that *F. novicida* dampens GBP recruitment by means of its atypical bacterial envelope. Furthermore, while GBP1 recruitment has been studied with *S. flexneri and S. typhimurium,* GBP1 recruitment to *F. novicida* appears to be less resilient to mutations, allowing us to highlight multiple independent features required for GBP1 targeting to bacteria.

Four features in GBP1 drive recruitment to *F. novicida*. Two of them, the GBP1 CAAX box or the C-terminal triple arginine patch also promote recruitment to *S. flexneri* and *S. typhimurium* (7, 15, 20). Interestingly, the two others were required for GBP1 targeting to *F. novicida* but were fully facultative for *S. flexneri.* They consist in a lysine residue (K_63_) present in a patch of three consecutive positively charged residues (KKK_61-63_) in the N-terminal region of GBP1 and a G_68_, the mutation of which ablates GBP1’s GDPase activity while leaving its GTPase activity intact (43). GMP production and its ensuing catabolism in uric acid contributes to NLRP3 inflammasome activation during *Chlamydia trachomatis* infection (43). Depending on the infecting pathogen, the GDPase activation may thus have two synergistic functions to promote inflammasome activation, the first one at the bacterial surface to assemble the caspase-4-activating platform (19, 20, 23) and the second one to activate the NLRP3 inflammasome. The role of GBP1 GDPase activity to target *F. novicida* was confirmed by replacing the guanine cap of GBP1 (which plays a specific role in GMP formation (11)) with the one of GBP2. Contrary to the G68A mutation, the guanine cap exchange decreased *S. flexneri* targeting. The guanine cap of GBP1 contains a third positive patch (RRK_243-245_) that is mirrored by a KKY_243-245_ in GBP2. Although, the RRK_243-245_ was not required for GBP1 binding to *E. coli* LPS in an in vitro assay (19), we cannot exclude that this patch may play a role in *S. flexneri* targeting possibly in synergy with the GDPase activity. Altogether, our study extends the findings from previous publications demonstrating that several independent GBP1 features contribute to targeting to Gram-negative bacteria. Furthermore, studying *F. novicida* targeting in comparison with *S. flexneri* identified specific GBP1 features that are required for *F. novicida* but facultative for *S. flexneri* targeting suggesting that GBP1 has evolved these different domains to recognize a diversity of pathogens.

In addition to GBP1 prenylation, we observed that GBP2 recruitment to *F. novicida* and *S. flexneri* was dependent on the CAAX box. This result indicates that further GBP specificities exist to control GBP recruitment downstream of GBP1. Indeed, GBP3 and GBP4, which are recruited to *S. flexneri* are devoid of a CAAX box while the prenylated GBP5 is not recruited. More surprisingly, the presence of the GBP2 CAAX box on either GBP1 or GBP2 was associated with a significantly higher recruitment to *F. novicida* than GBP1 or GBP2 with GBP1 CAAX box. GBP1 and GBP2 CAAX boxes drive farnesylation or geranylgeranylation, respectively (Table 1). The longer size of the GBP2 lipid anchor (20 carbons) might increase the stability of GBP recruitment in the membrane of *F. novicida*, which displays long lipid A acyl chains with 16 to 18 carbons (Fig. 7A). The factors controlling recruitment of GBPs downstream of GBP1 are still elusive. The current model states that pioneer prenylated GBPs recruit other GBPs through heterotypic interactions (1, 12). Both GBP2 and GBP5 interact with GBP1 in overexpression systems (8, 47) while GBP5 is not recruited on cytosolic bacteria indicating that GBP1 interactions are not the only drivers of GBP recruitment. Our chimera experiments (Fig. 4) mapped the GBP2 region directing recruitment to the central helical domain (K340-R535, spanning the second half of the α9 helix to the first half of the α12 helix). The knowledge on the function of this central GBP domain is still sparse (10, 26, 40). According to the current model, GBP1 polymerization requires the opening of a α9-α12 structural hairpin (26). The central helical domain of GBP2 identified may thus allow structural rearrangement to accommodate polymerization with GBP1, whereas this conformational change might not be possible in GBP5, at least on the bacterial surface. The structural requirements for GBP1 homopolymerization are now well known (24). Yet, GBP heteropolymer formation awaits to benefit from similar exquisite biochemical studies to provide an understanding of how the central helical domain identified here drives the specific recruitment of GBP2 to *F. novicida*.

In addition to the above discussed host features, our work further revealed that bacterial factors control GBP recruitment. Indeed, *F. novicida* escapes targeting by the non-prenylated GBP3 and GBP4. GBP2-4 recruitment to *S. flexneri* or *S. typhimurium* depend solely on GBP1 and on no other GBP (15, 19) suggesting that no further recruitment hierarchy exists downstream of GBP1. However, facing two different pathogens in the same experimental system, GBP2, GBP3 and GBP4 were differentially recruited – only GBP2 was targeted to *F. novicida*. Therefore, additional mechanisms control the selective GBP recruitment downstream of GBP1, and those mechanisms are dependent on bacterial factors. GBP3/4 might require the presence of a specific bacterial molecule that would act, together with GBP1, as a co-receptor to allow GBP3/4 recruitment, and that would be absent from *F. novicida* envelope. Alternatively, GBP1/2 polymer conformation at the surface of *F. novicida* may not be favorable to promote GBP3 or GBP4 binding. While the O-chain moiety of LPS drives GBP1 encapsulation of bacteria (20), several evidence suggest that other LPS domains contribute to GBP recruitment/function. First, GBPs are recruited at the surface of *S. flexneri* rough mutants (without O-chain) although at lower levels than to WT strain (21). Second, in vitro, GBP1 still binds *S. flexneri* Δ*rfaL* rough mutant although it does not promote GBP1 encapsulation (20). Third, mGBPs are required for full inflammasome activation in response to smooth LPS, rough LPS or even synthetic lipid A (48). Importantly, our work suggests that one of the bacterial factor that controls GBP recruitment is the number of lipid A acyl chains. Indeed, a clear and statistically significant enrichment of GBP3 was observed on *ΔlpxF F. novicida* mutant, which bears a penta-acylated lipid A. Interestingly, LPS from *ΔlpxF F. novicida* mutant is also recognized by caspase-11 (45) suggesting a convergent evolution of cytosolic LPS sensors. Of note *lpxF* deletion also results in the presence of an additional phosphate group in the disaccharide anchor of lipid A, which is absent in wild-type *F. novicida* but is otherwise present in enterobacteria LPS (Fig 7A). This phosphate addition increases the negative charge of the LPS and may thus account for or contribute to the recruitment of GBP3.

Interestingly, the GBP3 recruitment observed on *ΔlpxF* mutant strain was not comparable to the GBP1/2 recruitment observed on WT *F. novicida* or to the GBP3 recruitment observed on *S. flexneri*, suggesting that besides lipid A tetra-acylation, *F. novicida* has evolved other strategies to hide from GBP3 (and GBP4) recruitment. In addition to the tetra-acylation of its LPS, *F. novicida* has numerous other unique envelope characteristics, including a high proportion of free lipid A, and the presence of additional sugars in the lipid A anchor. Multiple properties of its bacterial envelope may thus cooperate to enable escape from GBP targeting. Finally, the highly virulent *F. tularensis* subspecies tularensis evades mGBP-mediated growth restriction more efficiently than *F. novicida* (36). Future studies should examine the role of the *F. tularensis* envelope “invisibility cloak” in the escape of GBP-mediated immune responses.

## MATERIALS AND METHODS

### Ethics statement

Blood from healthy donors was obtained from the Etablissement Français du Sang Auvergne-Rhône Alpes, France under the convention EFS 16-2066. Informed consent was obtained from all subjects in accordance with the declaration of Helsinki. Ethical approval was obtained from Comité de Protection des Personnes SUD-EST IV (L16-189).

### Bacterial strains

*Francisella novicida* strain Utah (U112) and related mutants were grown in Tryptic Soy agar and broth (Pronadisa) supplemented with 0.1% w/v L-cysteine. *F. novicida* ΔFPI and Δ*lpxF* mutants were previously described (49, 50). Lipid A composition of the Δ*lpxF* mutant was validated by mass spectrometry. *Shigella flexneri* str. M90T ΔipaH9.8 (51) was grown in Tryptic Soy agar and broth. *Escherichia coli* DH5α were grown in Lysogeny broth and agar (Pronadisa) supplemented with ampicillin (100 µg/ml) or kanamycin (30 µg/ml) when necessary.

### Cell cultures

U937 cells were maintained in RPMI Medium 1640 - GlutaMAX™-I (ThermoFisher Scientific) supplemented with 10% v/v fetal calf serum. HEK293T cells were maintained in Dulbecco’s Modified Eagle Medium with GlutaMAX™-I (ThermoFisher Scientific) with 10% v/v fetal calf serum and geneticin (200 µg/ml). Primary human CD14^+^ monocytes were isolated from blood and differentiated into macrophages for a week as previously described (33).

### Plasmid constructions

*GBP* fragments were amplified from pAIP plasmids containing *GBP1, GBP2, GBP5* cDNA in frame with a N-terminal HA tag-coding sequence using primers listed in Table S1. Chimeric *GBP* sequences were produced by joint PCR using a Phusion® High-Fidelity DNA polymerase (New England BioLabs) and purified from agarose gel using NucleoSpin® Gel and PCR clean up kit (MACHEREY-NAGEL). Point mutations were introduced into *GBP* sequences cloned in a pUC57 plasmid, using PFU Ultra II DNA polymerase. The methylated template was digested with DpnI and the mutated plasmid was transformed in *E. coli* DH5α cells through heat-shock transformation. The chimeric or mutated *GBP* sequences were then transferred into pAIP using restriction enzymes NotI and BamHI and the T4 DNA ligase (New England BioLabs). Transformed clones in *E. coli* DH5α were selected with ampicillin (100 µg/ml) for pAIP or kanamycin (30ug/ml) for pUC57 derivatives. All final constructs were verified by sequencing (Eurofins Genomics).

### Lentiviral production and generation of stable U937 cell lines

HEK293T cells were seeded in complete DMEM medium at 2.10^6^ cells per 25 cm^2^ cell culture flask and transfected 24h later with 4.3 µg pPAX2 (gag-pol expression), 1.43 µg pMDG (VSV-G expression) and 5.6 µg pAIP construct. Transfection was carried out in 1.4 ml OptiMEM^TM^ reduced serum medium (ThermoFisher Scientific) supplemented with 20 µM polyethylenimine (Sigma-Aldrich #408727). Complete DMEM was added 4 h later. On the following day, the medium was changed for 1.4 ml DMEM. Lentiviruses were collected 48 h after transfection. U937 cells were seeded at 2.5×10^5^ per well in a P24 plate and transduced by adding 500µl lentiviral particles to the culture medium. Starting from 72 h post-transduction, transduced cells were selected by treating with puromycin (2 µg/ml) for 14 days. Protein expression was controlled by Western Blot. All produced cell lines are described in Table S2. Control, *GBP1*^KO^, *GBP2*^KO^, *GBP3*^KO^ cells were previously described (23). *GBP4*^KO^ and *GBP5*^KO^ U937 cell lines were generated similarly using the following sgRNA: GBP4: GTAACCCTAAGAATGACTCG (guide 1), TGTGCGGTATAGCCCTACAA (guide 2); GBP5: AAACTCACCCGACCTTGACA (guide 1), GTTCACAGTATTGTACACAA (guide 2). Wild-type U937 and HEK-293T cells tested negative for *Mycoplasma*.

### Infections

To obtain macrophages, U937 monocytes were seeded 36h prior to infection in complete RPMI supplemented with 100 ng/ml phorbol myristate acetate (PMA, Sigma-Aldrich). Primary human monocytes were treated with 50 ng/ml M-CSF (Sigma-Aldrich) in complete RPMI for a week before infection. Treatment with 10^3^ U/ml hIFN-γ (Sigma-Aldrich) was done 18 h prior to infection unless otherwise specified. Bacteria were grown in overnight culture in 2ml TSB + 0,1% cysteine (*F. novicida*) or TSB (*S. flexneri*). Infection with *F. novicida* was done using overnight culture. For infection with *S. flexneri*, the overnight culture was diluted at 1/100 and the subculture was grown until it reached an OD_600nm_ 1. The bacteria were suspended in RPMI at the desired MOI and added onto the cells followed by a spinoculation at 1000g for 15 min (32°C). After 1 h of incubation at 37°C, the cells were washed and the medium was replaced in RPMI with gentamycin (5µg/ml for *F. novicida* or 100 µg/ml for *S. flexneri*) until the desired time post-inoculation. When applicable, chloramphenicol (4 µg/mL) was used to inhibit protein neosynthesis at 5 h p.i.

### Immunofluorescence

U937 macrophages and hMDMs were differentiated as described above and seeded at 5.10^5^ cells/ml in P12 plates (U937) or onto sterile glass coverslips (hMDMs). Infection was carried out as described above. At the indicated time of infection, the cells were washed and fixed with 2% formaldehyde (Sigma-Aldrich) in PBS for 10 min at RT. U937 cells were mounted on poly-L-lysine slides (Sigma-Aldrich) using Shandon Cytospin 3 cytocentrifuge for 10^5^ cells per slide. hMDMs were stained directly on the glass coverslips. Permeabilization was done with in PBS-Triton 0.1% for 10 min at RT. The samples were submerged in blocking buffer (5% BSA, 0.1% Triton, 0.02% NaN_3_ in PBS) for 1 h at RT or 4°C O/N then stained with the appropriate antibodies, see Table S3. DAPI (4′,6-diamidino-2-phenylindole at 100 ng/ml, ThermoFischer Scientific) was used for DNA staining. Coverslips were mounted using Fluoromount G^TM^ (Invitrogen) mounting medium. Images for statistical analysis were taken on a Nikon Eclipse Ts2R-FL inverted microscope. For representative images, the samples were imaged on a Zeiss LSM 800 confocal microscope or Zeiss Elyra 7 SIM/STORM microscope. ImageJ software was used for analysis.

### Phagosomal rupture assay

Quantification of vacuolar *F. novicida* escape was done using the β-lactamase/CCF4 assay (Life Technologies)(52). Briefly, U937 monocytes were seeded in P48 plates at 1.25×10^5^ cells/well in PMA-supplemented RPMI. The cells were treated with 10^3^ U/lm IFNγ and infected as described above. Three h post-infection, the cells were washed and incubated in CCF4 for 1 h at RT in the presence of 2.5 mM probenecid (Sigma-Aldrich). Live cells (propidium-iodide negative, CCF4 positive) were tested for *F. novicida*-mediated phagosomal rupture by flow cytometry using excitation at 405 nm and detection at 450 nm (cleaved CCF4) or 510 nm (intact CCF4).

### Cell death assay

U937 cells were differentiated for 10^5^ cells/well in 96 well plates, treated with 100 U/ml IFNγ and infected with *F. novicida* at MOI 100 as described above. One h p.i. the cells were washed with PBS and the medium was replaced with CO2-independent medium supplemented with 10% FCS, 5µg/ml gentamycin and 5µg/ml propidium iodide (ThermoFischer Scientific). PI fluorescence was measured every 15 min during 24 h on a Tecan microplate fluorimeter. Data were normalized using uninfected cells and cells treated with 1% Triton X100 (100% cell death).

### Real time PCR

PMA-differentiated U937 were treated or not with IFNγ and infected or not as described above. Total RNA was extracted using chloroform and TRI Reagent ® (Sigma-Aldrich #93289) and reverse transcribed with random primer combined with Im-Prom Reverse Transcription System (Promega #A3800). Quantitative real-time PCR was performed using FastStart Universal SYBR Green Master Mix (Roche #04913850001) and an Applied StepOnePlus^TM^ Real-Time PCR System (ThermoFisher Scientific). Gene-specific transcript levels were normalized to the amount of human *HPRT* transcripts. Primer sequences are available in (33).

### Western blotting

U937 cells were washed in PBS and lysed for 30 min on ice using Radioimmunoprecipitation buffer supplemented with EDTA-free protease inhibitor cocktail cOmplete^TM^ (Roche). Cleared lysate was obtained by centrifugation at 11 000g for 10 min at 4°C. Total protein concentration of the lysates was determined using a Micro BCA^TM^ Protein Assay Kit (ThermoFisher Scientific) according to the manufacturer’s protocol. Laemmli Sample Buffer 4X (Bio-Rad) with 10% v/v β-mercaptoethanol (Sigma-Aldrich) was added to the protein samples before boiling them for 10 minutes at 95°C. Protein extracts were deposited onto a 4-15% Mini-PROTEAN® TGX^TM^ precast protein gel (Bio-Rad) for migration. Following migration, the samples were transferred onto a membrane using Trans-Blot® Turbo^TM^ RTA transfer system and kit (Bio-Rad). The membranes were saturated with 5% skimmed milk, then stained with the appropriate primary and secondary antibodies (Table S3) and revealed with an ECL Western Blotting detection reagent (Dd Biolab).

### Statistical analysis

Statistical analysis was performed with GraphPad Prism 9 software. Normality was assessed for data sets with n > 20 entries using D’Agostino & Peerson omnibus normality test. Multiple comparison was done with analysis of variance tests with post-hoc corrections (Dunnett’s or Sidak’s depending on the selected comparisons). In the case of single comparisons, two tailed t tests were performed.

### Data availability

All relevant data are available in the paper or the supplementary material. Plasmid constructs have been deposited in Addgene (public access pending). Additional data is available upon request.

## ACKNOWLEDGMENTS

We thank D. Monack (Stanford University) and A. Cimarelli (CIRI) for reagents. We acknowledge the contribution of SFR Biosciences (UAR3444/CNRS, US8/Inserm, ENS de Lyon, UCBL) flow cytometry and imaging facilities (especially Elodie Chartre). This project is supported by an ANR grant to TH (Tulamibe, ANR-17-ASTR-0024-02). SVV is supported by a scholarship from AID (AID 2019013) & Inserm.

---

**FIG S1.**
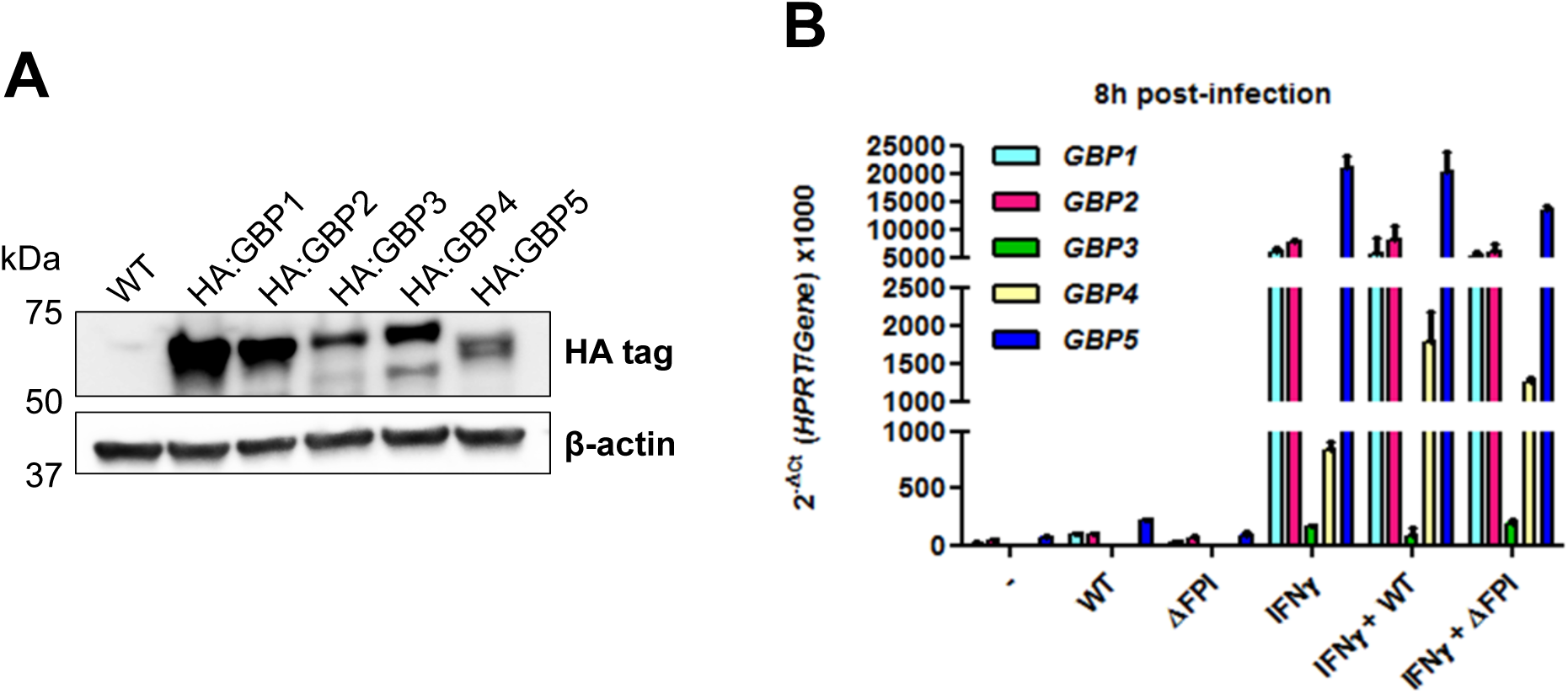
(A) Stable expression of HA-tagged GBPs was assessed by Western blot in U937 cell lines. (B) Endogenous levels of *GBP* transcripts were quantified by qRT-PCR and normalized to *HPRT* transcript levels after treatment with IFNγ or infection with WT or ΔFPI *F. novicida* strains.

**FIG S2.**
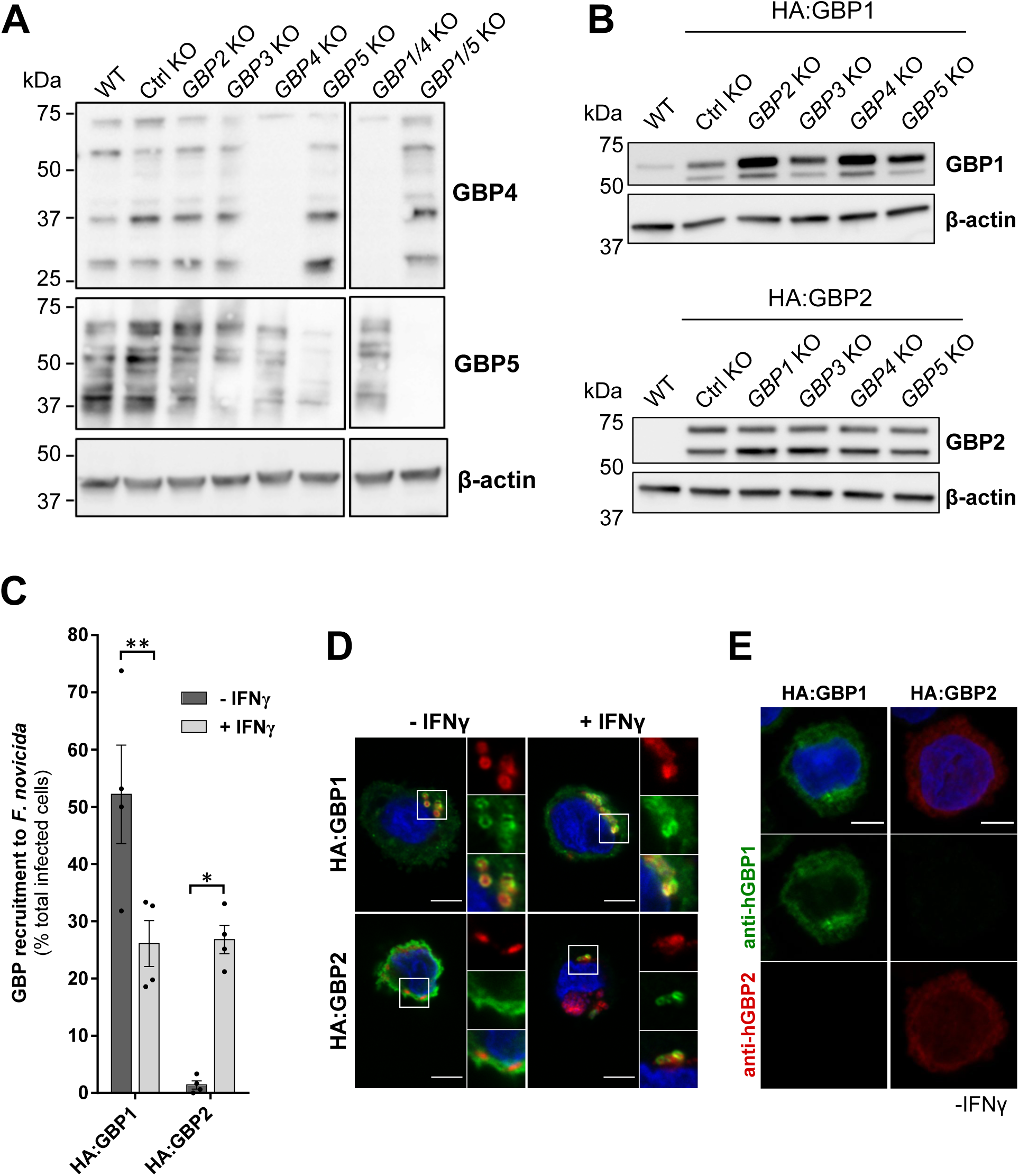
(A) *GBP* or control KO U937 cells were assessed for GBP4 or GBP5 expression by Western blot. The residual GBP5 signal in *GBP5*^KO^ is due to cross-reactivity of the antibody with GBP1 as demonstrated with *GBP1/5^DKO^*. (B) Stable expression of HA:GBP1 or HA-GBP2 were analyzed by Western blot in the indicated U937 cell lines. (C) GBP recruitment was scored as the percentage of infected cells with GBP-bacteria colocalization in U937 macrophages in the presence or absence of IFNγ. ANOVA with Sidak’s multiple analysis test was used: **, p < 0.01. (D) Representative images with scale bar 5µm and 3X zoom are shown. (E) Specificity of anti-hGBP1 and anti-hGBP2 antibodies was illustrated with confocal images acquired with identical imaging settings for both samples. The images are shown without brightness and contrast adjustments. Scale bar, 5 µm.

**FIG S3.**
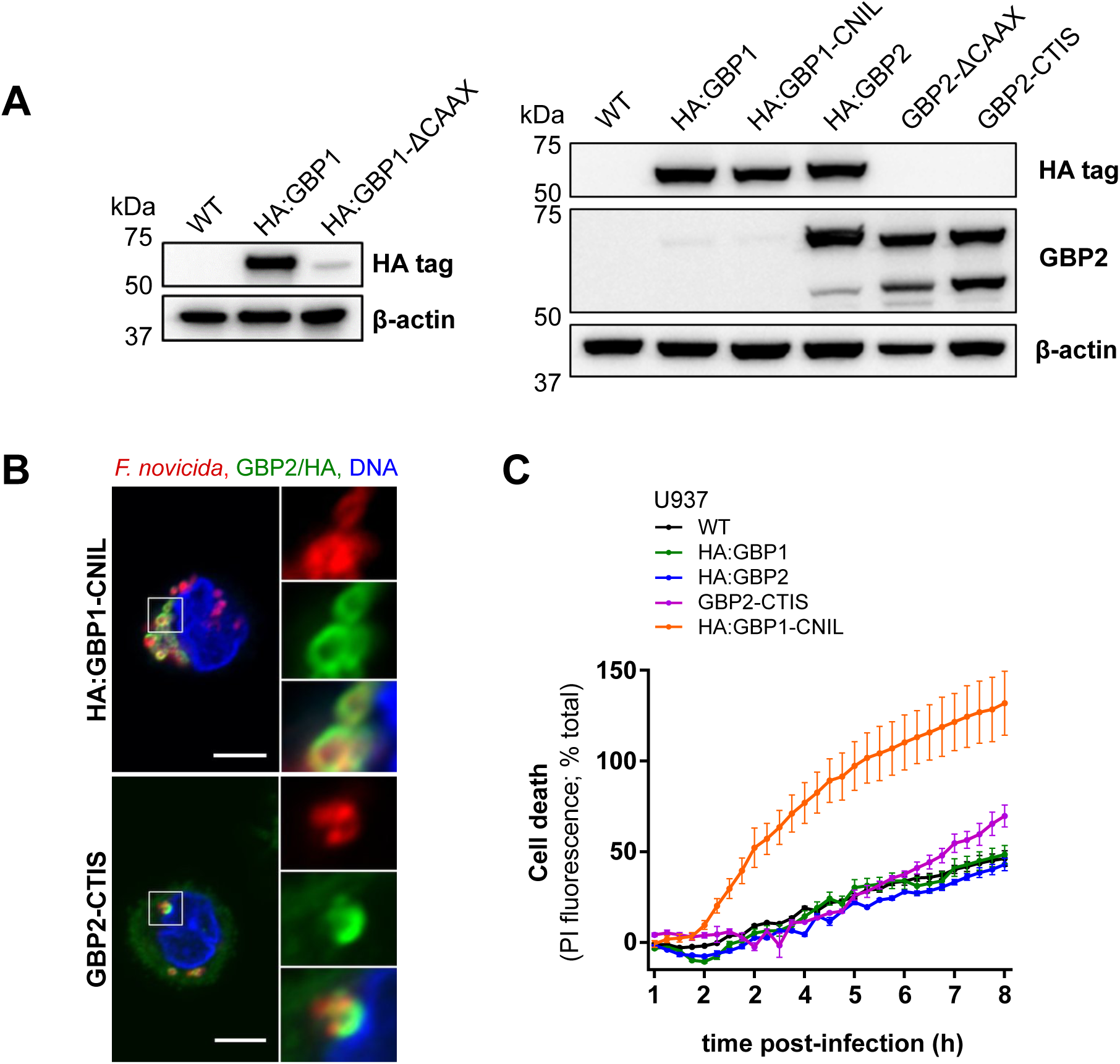
(A) Stable expression of the indicated GBP constructs was analyzed by Western blot in U937 cell lines. (B) Representative images of IFNγ-treated U937 macrophages, infected with *F. novicida* for 7 h are shown. Scale bar, 5 µm with 2X zoom on the right panels. (C) Propidium iodide (PI) incorporation/fluorescence was monitored every 15 min in IFNγ-primed macrophages, infected with *F. novicida* at MOI 100 and normalized to untreated cells and to Triton X100-treated cells. Each point corresponds to the mean +/- SEM of a biological triplicate from one experiment representative of 3 independent experiments.

**FIG S4.**
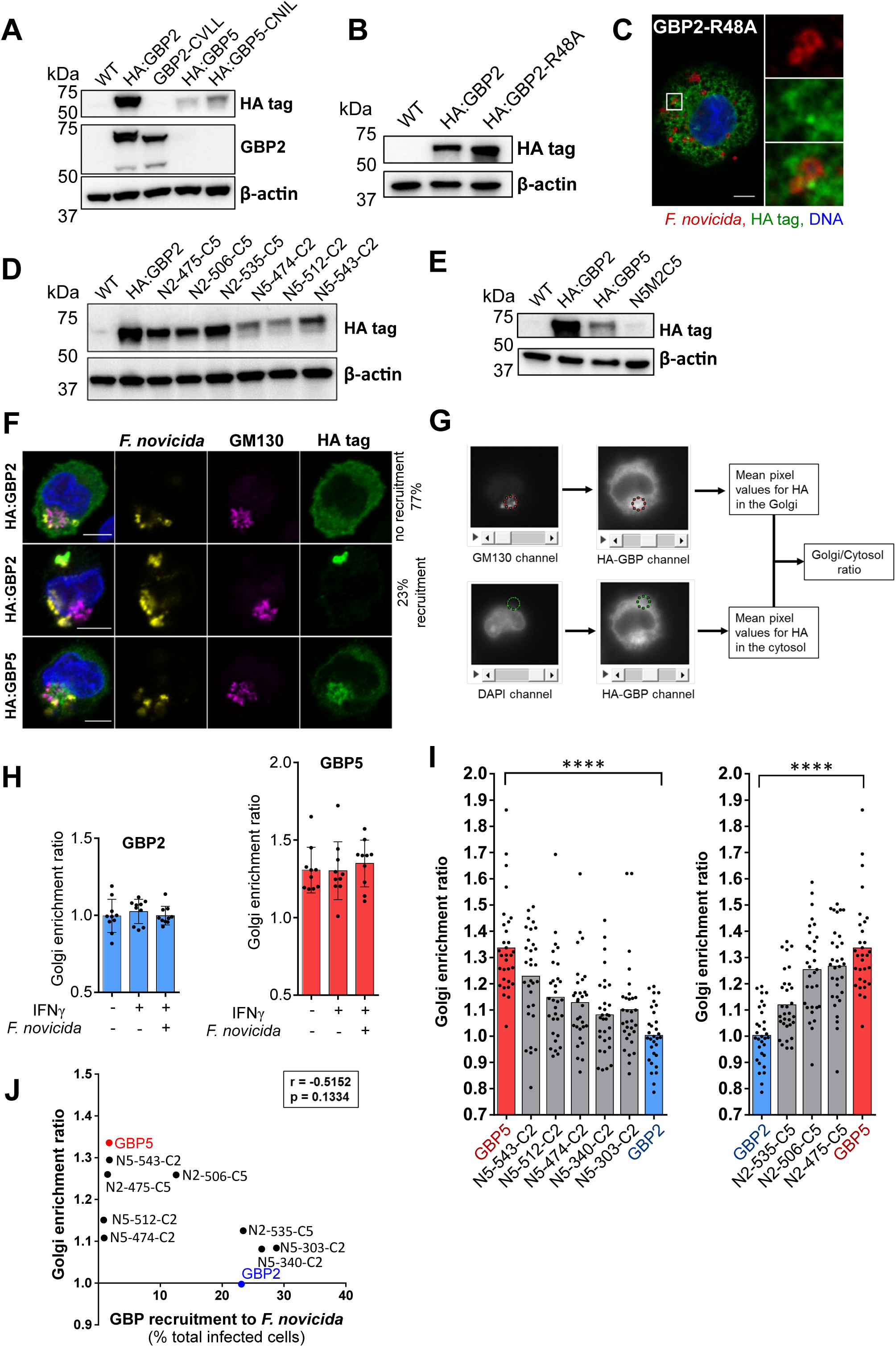
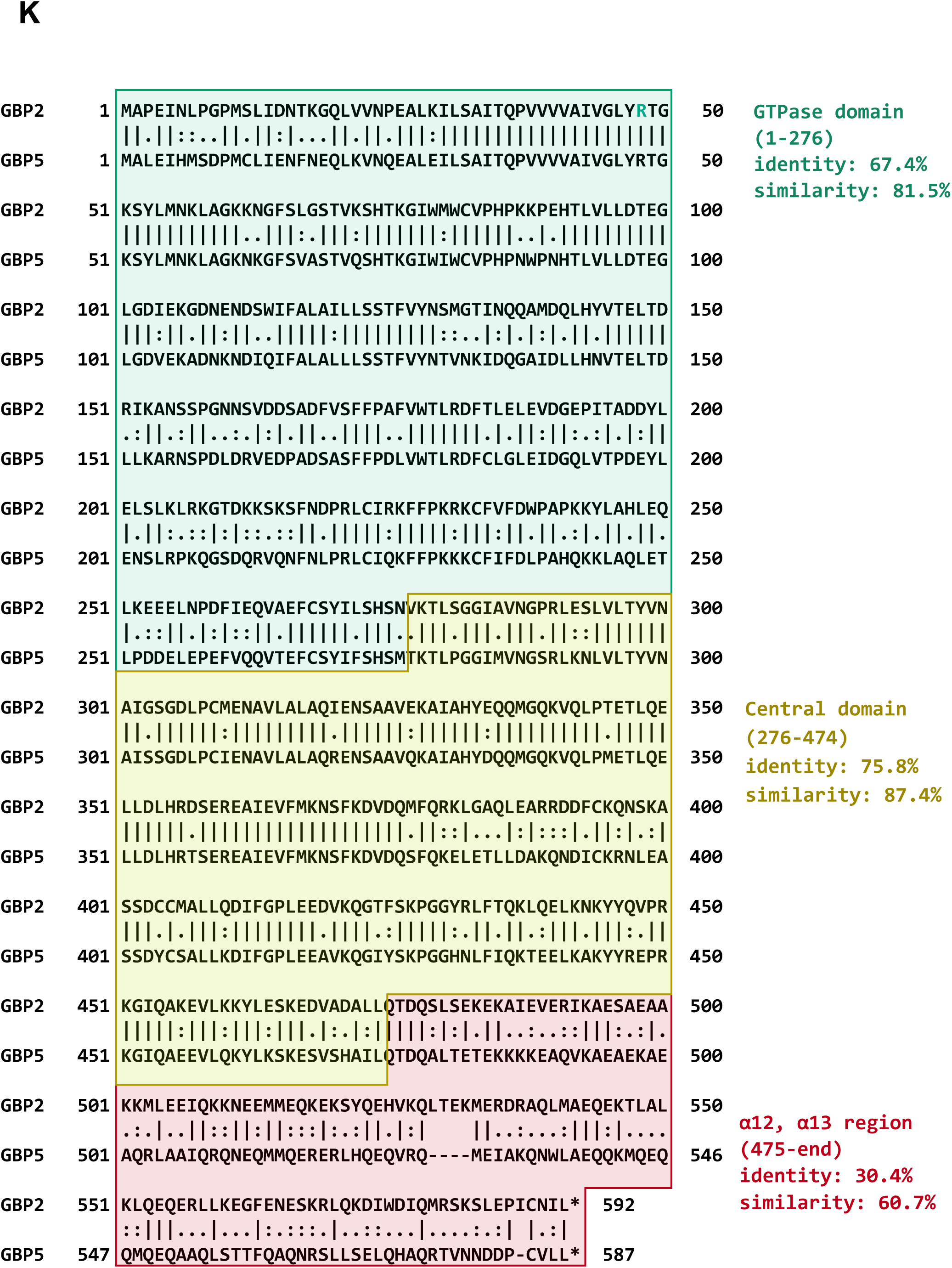
(A, B, D, E) Stable expression of HA-tagged GBP constructs was analyzed by Western blot in U937 cells. (C, F, H) Representative images of IFN-γ-treated, U937 macrophages infected with *F. novicida* are presented with a 5 µm scale bar. (G) Golgi enrichment was scored using GM130 staining to delineate the Golgi region and extract the corresponding pixel intensity values for the HA-GBP image channel. A similar area outside of the nucleus served to obtain a cytosol baseline value. (H, I) Golgi enrichment ratios of infected (H) or untreated (H, I) U937 cells. Each point represents the Golgi enrichment ratio calculated for a single cell. The bar represents the mean of three independent experiments with the Golgi enrichment ratio calculated in 10 or more cells per experiment. Statistical differences between GBP2 and GBP5 (****, p < 0.0001) were evaluated through two tailed Mann-Whitney analysis. (J) The mean Golgi enrichment of each construct is plotted in function of the mean GBP recruitment. Spearman’s correlation results are presented. (K) GBP2 and GBP5 protein sequences were aligned with Emboss-Needle Pairwise Sequence Alignment using the BLOSUM 62 matrix. Local identity and similarity were evaluated by separate alignments with the same matrix.

**FIG S5.**
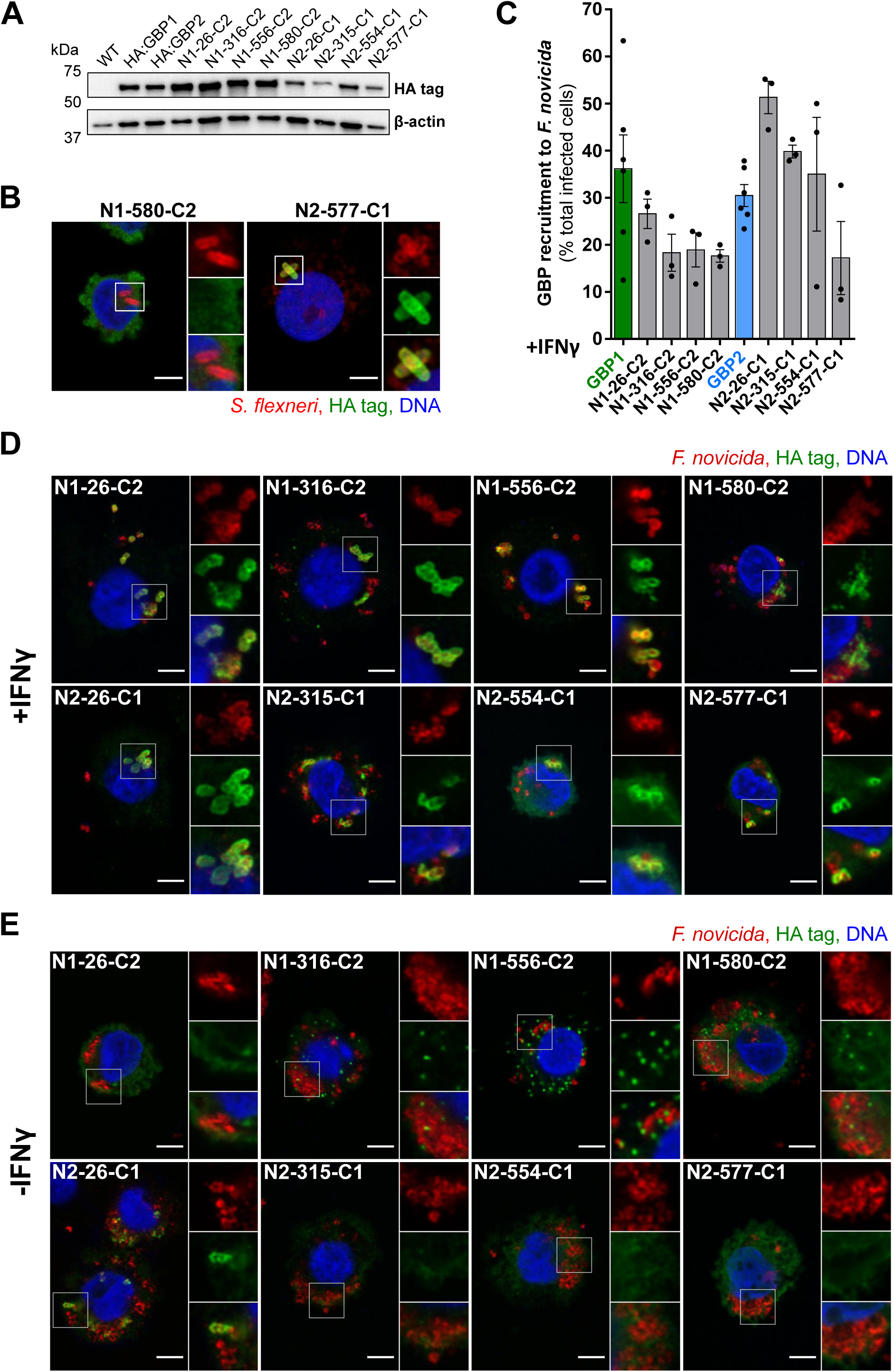
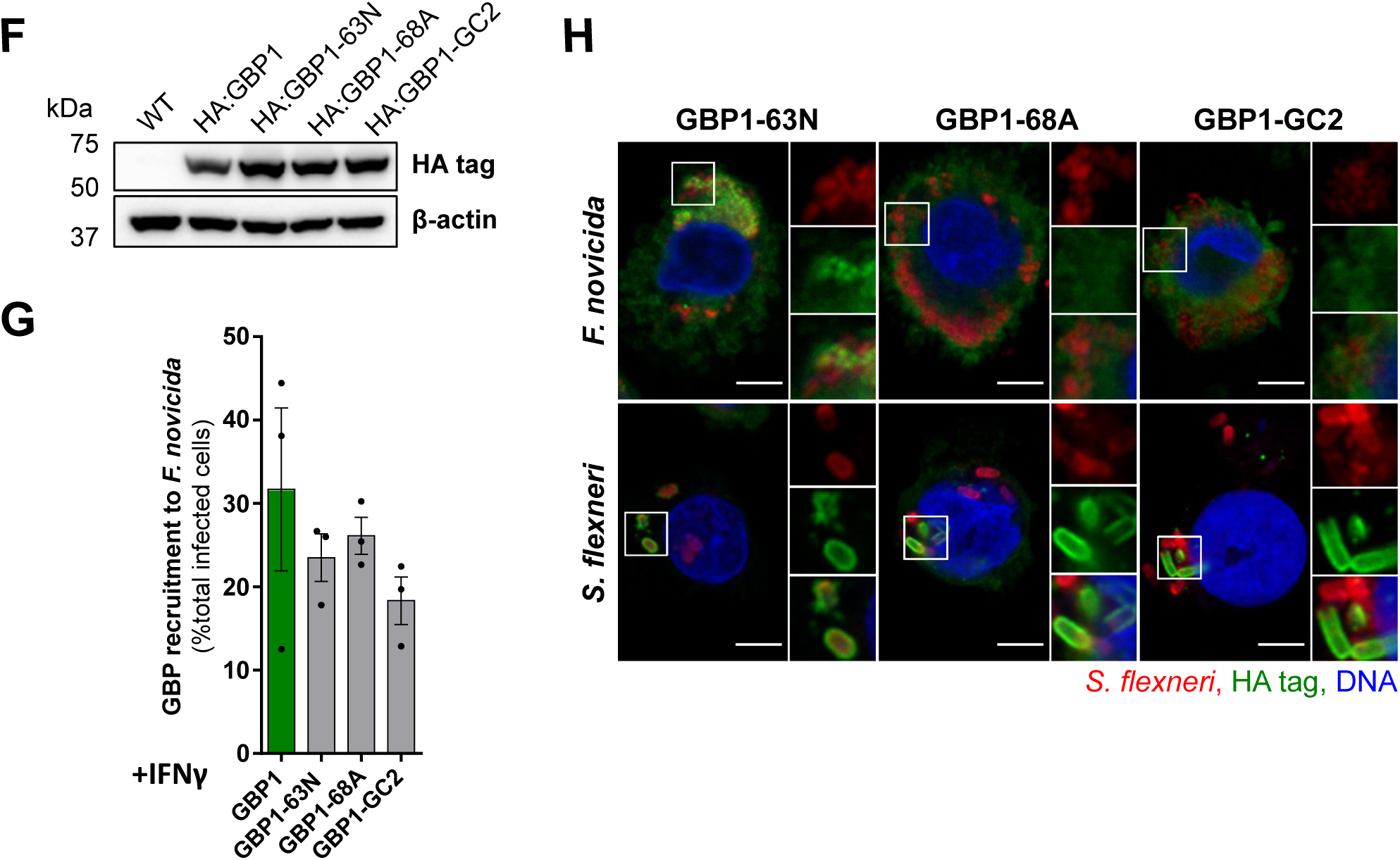
(A, F) Stable HA-tagged GBP expression in U937 cells was analyzed by Western blot. IFNγ-primed (B-D, G) or untreated (E, H) U937 macrophages were infected with *F. novicida* for 7 h (C-E, G) or *S. flexneri* ΔipaH9.8 for 90 min (B, H). (B, D, E, H) Representative images are shown with a scale bar of 5µm and a 2X zoom. (C, G) GBP recruitment was quantified as the percentage of infected cells with HA-GBP-bacteria colocalization. Each point corresponds to the value from one experiment with 50-100 infected cells analyzed. The bar represents the mean +/- SEM of three to six independent experiments. ANOVA with Sidak’s analysis did not demonstrate statistical differences in recruitment between GBP chimeric/mutated and control cell lines.

**FIG S6.**
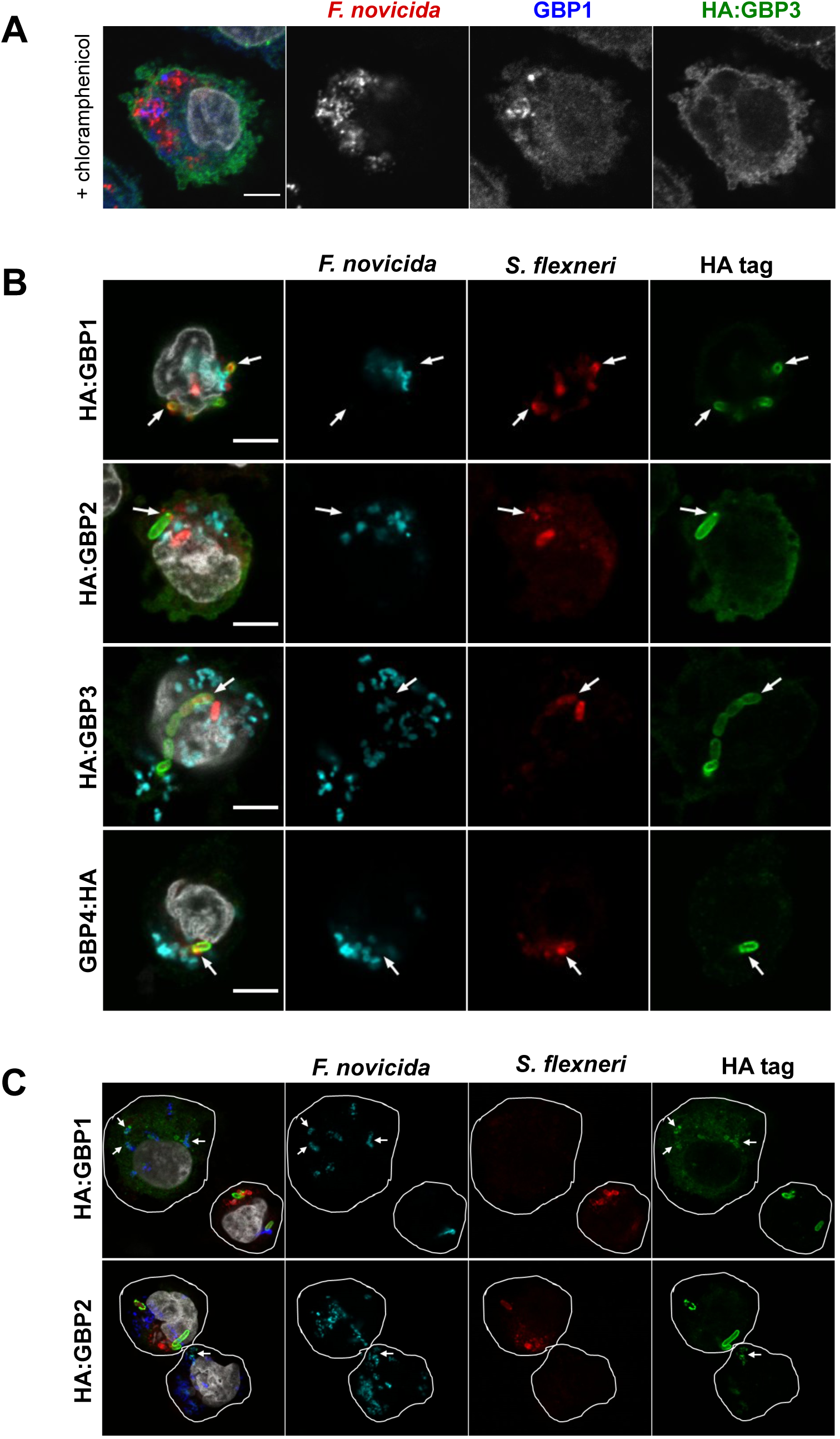
(A) *F. novicida*-infected U937 macrophages were treated with chloramphenicol for 2 h at 5 h p.i. Representative images are shown. Scale bar, 5µm. (B) IFNγ-primed U937 macrophages were co-infected with *F. novicida* and *S. flexneri* ΔipaH9.8 for 4 h. Representative images are shown. The arrows show GBP recruitment to *S. flexneri*. (C) GBP1 and GBP2 are recruited to *F. novicida* (arrows) in cells neighboring co-infected cells. The arrows show HA-GBP recruitment to *F. novicida*. Cell perimeter (white line) was delimited using a mask on HA staining as a cytosolic marker.

**FIG S7.**
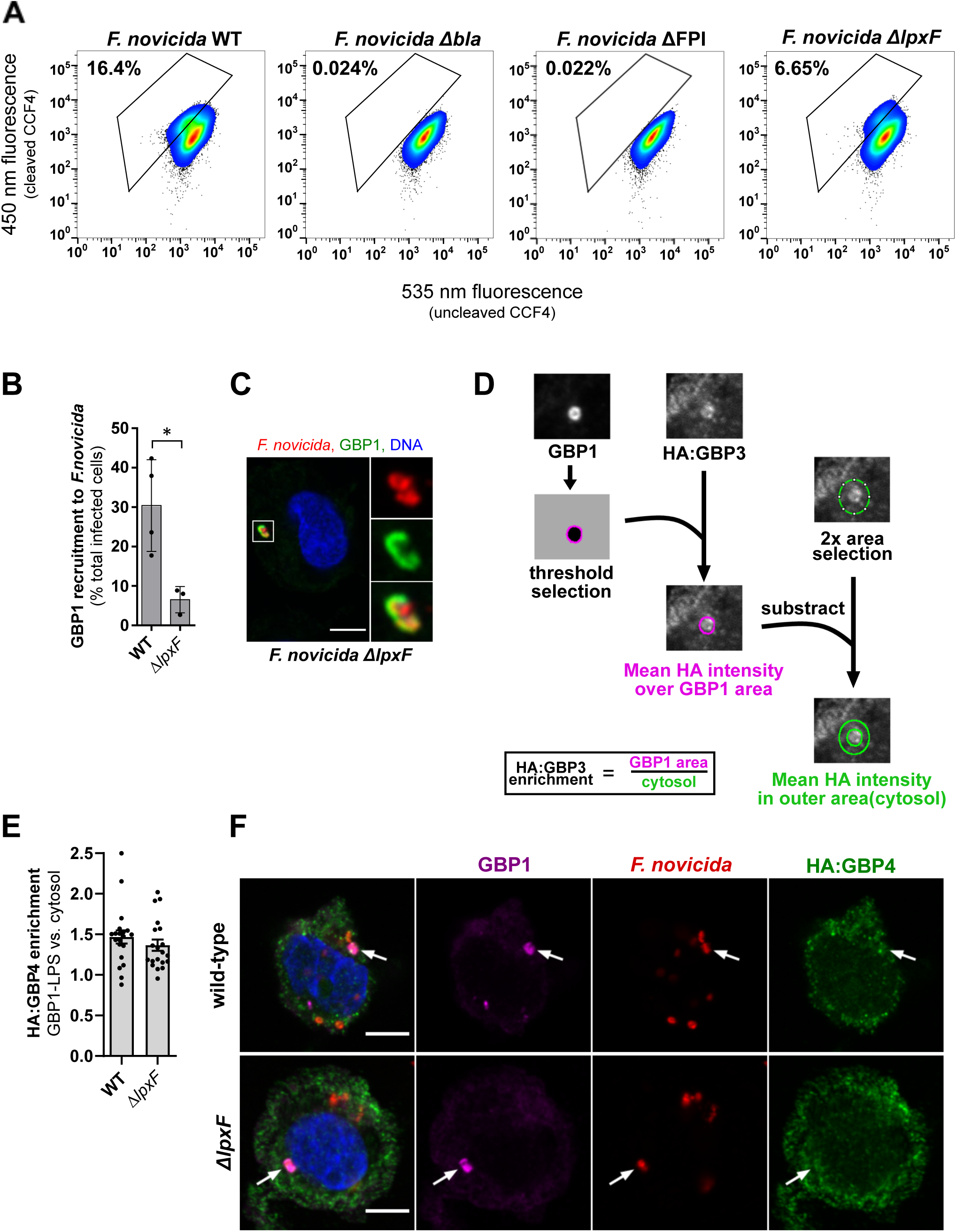
IFNγ-treated U937 macrophages were infected with *F. novicida* for 7 h. (A) Phagosomal rupture in *F. novicida*-infected macrophages, was evaluated by CCF4/β-lactamase flow cytometry assay. The cytosolic β-lactam FRET probe CCF4 emits at 535 nm when uncleaved, and 450 nm when cleaved by *F. novicida* β-lactamase. Mutants in the β-lactamase gene (*Δbla*) or in the Francisella Pathogenicity Island (ΔFPI) are presented as controls. (B) HA:GBP recruitment was calculated as the percentage of infected cells with HA-GBP-*F. novicida* colocalization. Two-tailed t test with Welch’s correction: *, p < 0.05. (C) Representative image of endogenous GBP1 recruitment to *F. novicida* Δ*lpxF* strain is shown. Scale bar 5 µm with 3X zoom. (D) Pipeline for scoring HA:GBP3 (or HA:GBP4) enrichment on one bacterium (or a cluster of bacteria) targeted by endogenous GBP1. (E) Each point represents the HA:GBP4 enrichment value of a single GBP1 recruitment area. Two-tailed Mann-Whitney analysis did not reveal any statistical difference in HA-GBP4 recruitment between WT and Δ*lpxF* strains. (F) Representative images are shown, scale bar 5 µm.

**Table S1.**
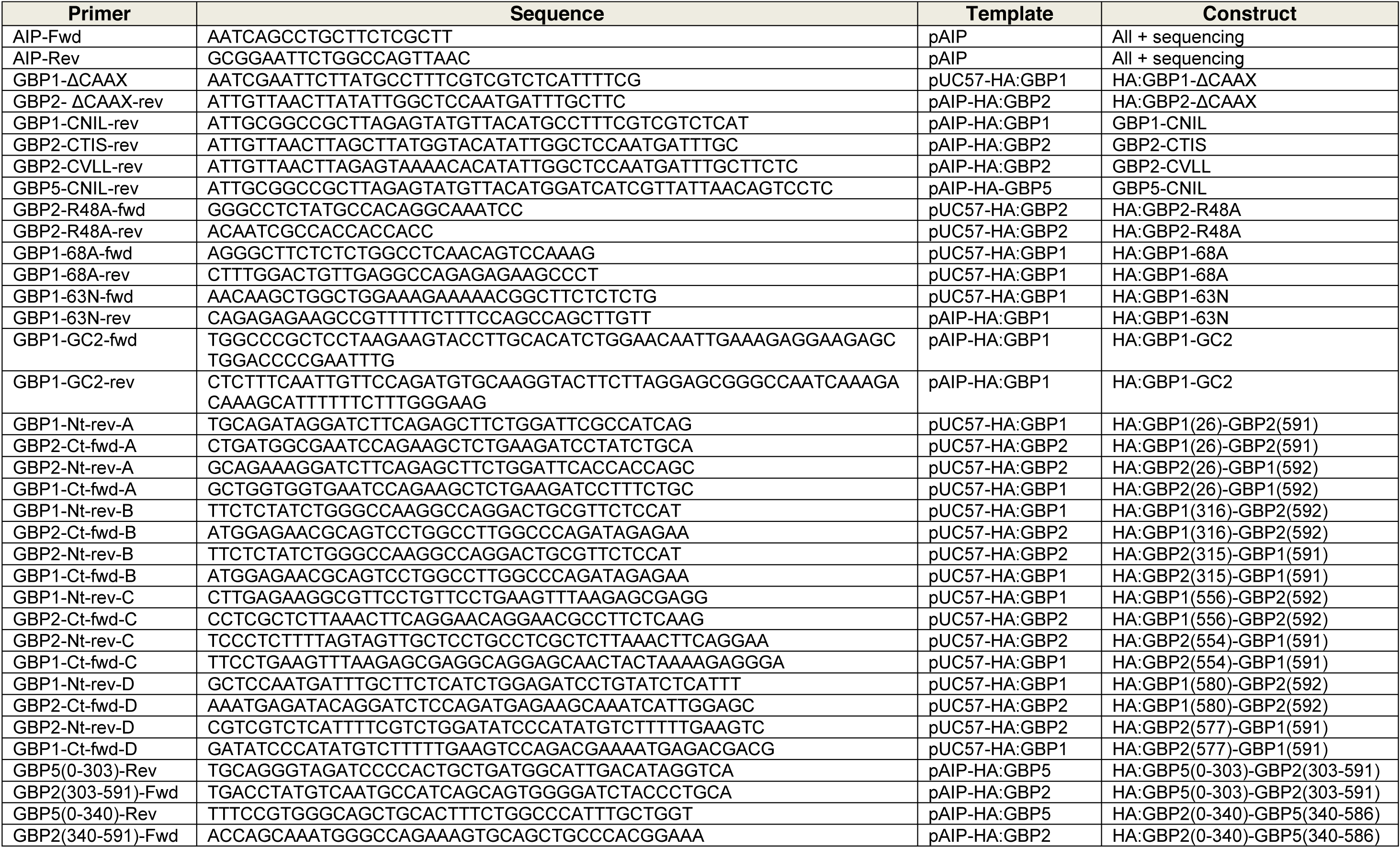
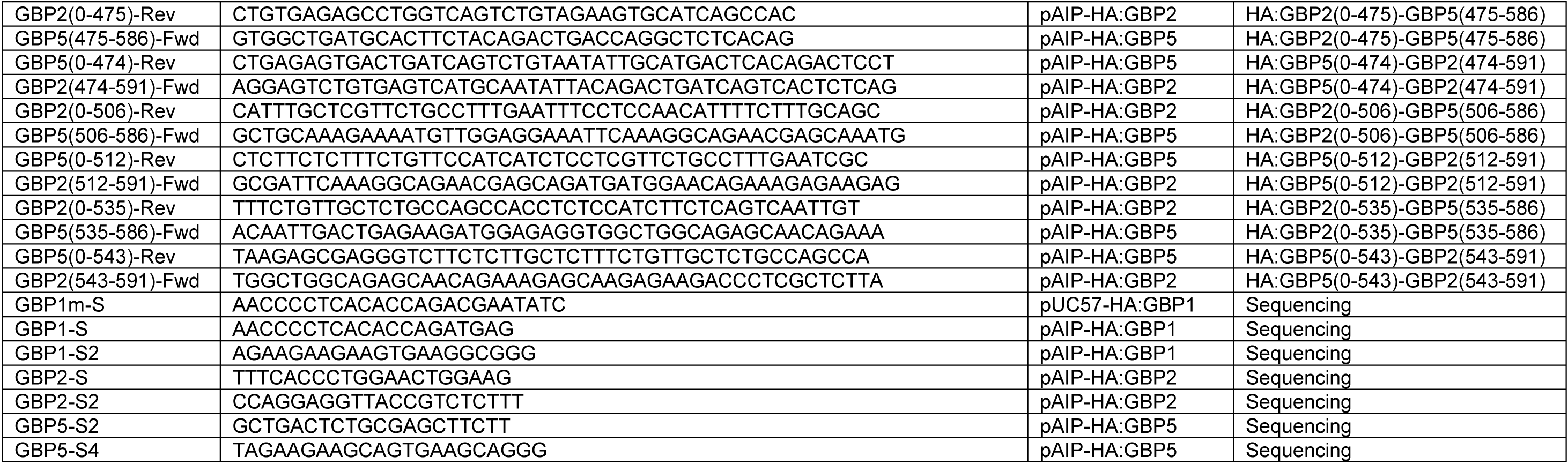
Primers

**Table S2.** Cell lines

*Note:* All U937 GBP cell lines are transduced for a stable expression of a HA-tagged construct unless otherwise specified.
| Cell line | Background | Description | Reference |
| --- | --- | --- | --- |
| HA:GBP1 | U937 WT | Stable expression; * | Santos *et al.*, 2018 (48) |
| HA:GBP2 | U937 WT | Stable expression; * | Santos *et al.*, 2018 (48) |
| HA:GBP3 | U937 WT | Stable expression; * | This study |
| HA:GBP4 | U937 WT | Stable expression; * | This study |
| HA:GBP5 | U937 WT | Stable expression; * | Santos *et al.*, 2018 (48) |
| OR5B17 KO | U937 WT | Crispr-Cas9 KO (control) | Wandel *et al.*, 2020 (23) |
| GBP1 KO | U937 WT | Crispr-Cas9 KO | Wandel *et al.*, 2020 (23) |
| GBP2 KO | U937 WT | Crispr-Cas9 KO | Wandel *et al.*, 2020 (23) |
| GBP3 KO | U937 WT | Crispr-Cas9 KO | Wandel *et al.*, 2020 (23) |
| GBP4 KO | U937 WT | Crispr-Cas9 KO | This study |
| GBP5 KO | U937 WT | Crispr-Cas9 KO | This study |
| GBP1/5 KO | U937 GBP1 KO | Crispr-Cas9 KO | This study |
| GBP1/4 KO | U937 GBP1 KO | Crispr-Cas9 KO | This study |
| OR5B17 KO<br>HA:GBP1 | U937 OR5B17<br>KO | Stable expression | This study |
| GBP2 KO HA:GBP1 | U937 GBP2 KO | Stable expression | This study |
| GBP3 KO HA:GBP1 | U937 GBP3 KO | Stable expression | This study |
| GBP4 KO HA:GBP1 | U937 GBP4 KO | Stable expression | This study |
| GBP5 KO HA:GBP1 | U937 GBP5 KO | Stable expression | This study |
| OR5B17 KO<br>HA:GBP2 | U937 OR5B17<br>KO | Stable expression | This study |
| GBP1 KO HA:GBP2 | U937 GBP1 KO | Stable expression | This study |
| GBP3 KO HA:GBP2 | U937 GBP3 KO | Stable expression | This study |
| GBP4 KO HA:GBP2 | U937 GBP4 KO | Stable expression | This study |
| GBP5 KO HA:GBP2 | U937 GBP5 KO | Stable expression | This study |
| HA:GBP1-ΔCAAX | U937 WT | CAAX box deletion by simple PCR | This study |
| GBP2-ΔCAAX | U937 GBP2 KO | CAAX box deletion by simple PCR; no tag; * | This study |
| GBP2 | U937 GBP2 KO | KO complementation | This study |
| HA:GBP2-R48A | U937 WT | GTPase inactive, point mutation PCR | This study |
| HA:GBP1-CNIL | U937 WT | GBP2 CAAX box, simple PCR; * | This study |
| GBP2-CTIS | U937 GBP2 KO | GBP1 CAAX box, simple PCR; no tag; * | This study |
| GBP2-CVLL | U937 GBP2 KO | GBP5 CAAX box, simple PCR; no tag; * | This study |
| HA:GBP5-CNIL | U937 WT | GBP2 CAAX box, simple PCR; * | This study |
| HA:N1-26-C2 | U937 WT | GBP1 Nter – GBP2 Cter chimera, joint PCR | This study |
| HA:N1-316-C2 | U937 WT | GBP1 Nter – GBP2 Cter chimera, joint PCR | This study |
| HA:N1-554-C2 | U937 WT | GBP1 Nter – GBP2 Cter chimera, joint PCR | This study |
| HA:N1-580-C2 | U937 WT | GBP1 Nter – GBP2 Cter chimera, joint PCR; * | This study |
| HA:N2-26-C1 | U937 WT | GBP2 Nter – GBP1 Cter chimera, joint PCR | This study |
| HA:N2-315-C1 | U937 WT | GBP2 Nter – GBP1 Cter chimera, joint PCR | This study |
| HA:N2-556-C1 | U937 WT | GBP2 Nter – GBP1 Cter chimera, joint PCR | This study |
| HA:N2-577-C1 | U937 WT | GBP2 Nter – GBP1 Cter chimera, joint PCR; * | This study |
| HA:GBP1-63N | U937 WT | 63K to 63N, K patch; point mutation PCR; * | This study |
| HA:GBP1-68A | U937 WT | 68G to 68A, GDP hydrolase null; point mutation PCR; * | This study |
| HA:GBP1-GC2 | U937 WT | Guanine cap of GBP2, joint PCR; * | This study |
| HA:N2-475-C5 | U937 WT | GBP2 Nter – GBP5 Cter chimera, joint PCR | This study |
| HA:N2-506-C5 | U937 WT | GBP2 Nter – GBP5 Cter chimera, joint PCR | This study |
| HA:N2-535-C5 | U937 WT | GBP2 Nter – GBP5 Cter chimera, joint PCR | This study |
| HA:N5-303-C2 | U937 WT | GBP5 Nter – GBP2 Cter chimera, joint PCR | This study |
| HA:N5-340-C2 | U937 WT | GBP5 Nter – GBP2 Cter chimera, joint PCR | This study |
| HA:N5-474-C2 | U937 WT | GBP5 Nter – GBP2 Cter chimera, joint PCR | This study |
| HA:N5-512-C2 | U937 WT | GBP5 Nter – GBP2 Cter chimera, joint PCR | This study |
| HA:N5-543-C2 | U937 WT | GBP5 Nter – GBP2 Cter chimera, joint PCR | This study |
| HA:N5M2C5 | U937 WT | GBP2 (340-535) in GBP5 background, joint PCR | This study |

\* Negative *Mycoplasma* test (28.04.2021)

**Table S3.** Antibodies

Primary
| Antigen | Made in | Reference | Experiment (Fig.) | Dilution |
| --- | --- | --- | --- | --- |
| HA tag | mouse | Sigma-Aldrich #H3663 | **IF** (all except otherwise indicated) | 1:3000 |
| HA tag | rabbit | Cell Signaling #3724S | **WB** (all except S2.A, B) | 1:5000 |
| hGBP1 | rabbit | Abcam #ab131255 | **IF** (2.D, E, F; 7.B, C; S2. E; S6; S7. C, E, F), **WB** (S2.B) | IF 1:150;<br>WB 1:2000 |
| hGBP2 | mouse | Novus #NBP1-47768 | **IF** (2.D, E, F; 6.B; S2.B); non-tagged GBP2 mutants ; **WB** (S2.B) | IF 1:1000<br>WB 1:2000 |
| hGBP4 | rabbit | Prointech #17746-1-AP | **WB** (S2.A) | 1:1000 |
| hGBP5 | rabbit | Prointech #13220-1-AP | **WB** (S2.A) | 1:1000 |
| *F. tularensis* LPS | chicken | Denise Monack lab | **IF** (where indicated) | 1:1000 |
| *Shigella* group LPS | rabbit | Novus #NB100-65058 | **IF** (where indicated) | 1:500 |
| GM-130 | rabbit | Cell Signaling #12480 | **IF** (S4) | 1:3000 |
| actin | mouse | Sigma-Aldrich #A3853 | **WB** (all) | 1:5000 |

Secondary
| Anti-IgG | Conjugate | Reference | Experiment (Fig.) | Dilution |
| --- | --- | --- | --- | --- |
| chicken | Alexa Fluor 568 | Invitrogen #A11041 | **IF** (S1, S4) | 1:1000 |
| chicken | Alexa Fluor 594 | Life #A21207 | **IF** (all except otherwise indicated) | 1:1000 |
| chicken | Alexa Fluor 647 | Sigma-Aldrich #A21449 | **IF** (2.D, E, F; 6.A, B; 7.B, C; S6; S7.C, D ) | 1:1000 |
| mouse | Alexa Fluor 488 | Sigma-Aldrich #A10667 | **IF** (all where otherwise indicated) | 1:1000 |
| mouse | Alexa Fluor 594 | Sigma-Aldrich #A11032 | **IF** (Fig. 2D, E, F; S2. E) | 1:1000 |
| rabbit | Alexa Fluor 488 | Life #A21206 | **IF** (Fig. 2D, E, F; S2. E) | 1:1000 |
| rabbit | Alexa Fluor 594 | Invitrogen #A11012 | **IF** (all *S. flexneri* staining; 7.B, C; S6; 7.C,D; S7.C, E, F) | 1:1000 |
| rabbit | Alexa Fluor 647 | ThermoFisher #A21246 | **IF** (S4) | 1:500 |
| mouse | HRP | Promega #W402b | **WB** (all) | 1:5000 |
| rabbit | HRP | Sigma-Aldrich #A0545 | **WB** (all) | 1:5000 |

